# *Cryptosporidium* sequentially remodels its single rhoptry into the host interface

**DOI:** 10.64898/2026.07.21.739819

**Authors:** Allison Cohen, Tim Bergner, Amandine Guérin, Maryse Lebrun, Michael Laue, Christian Klotz, Boris Striepen

**Affiliations:** Department of Pathobiology, School of Veterinary Medicine, University of Pennsylvania, Philadelphia, PA, U.S.A.; Robert Koch Institute, Berlin, Germany; Department of Microbiology and Molecular Medicine, University of Geneva, Geneva, Switzerland; LPHI, CNRS, INSERM, Université de Montpellier, Montpellier, France

## Abstract

The apicomplexan parasite *Cryptosporidium* is a leading cause of diarrheal disease in young children. Within the intestinal epithelium, *Cryptosporidium* establishes a unique intracellular niche in the apical brush border of enterocytes. A structurally complex interface between host and parasite acts as a holdfast and enables protein and metabolite transport. How the parasite invades its host cell and builds the interface is poorly understood. Here, we reveal parasite invasion with high temporal and spatial resolution using rigorous molecular markers. We find that sequential discharge of specialized secretory organelles anchors the parasite within the host prior to internalization. The single *Cryptosporidium* rhoptry persists beyond its initial pre-invasion discharge, acting as a conduit for the secretion of multiple waves of effector proteins. Ultimately, the rhoptry membrane gives rise to the feeder organelle that separates host from parasite cytoplasm. These findings lead us to propose a mechanistic model of *Cryptosporidium* invasion and intracellular parasitism.

## Introduction

Infection with the parasite *Cryptosporidium* is a leading cause of diarrheal disease, which is most severe in individuals with primary or acquired immune deficiency and in young children[1]. Globally, Cryptosporidiosis is an important contributor to early childhood mortality, especially in resource-poor settings, and it is linked to malnutrition both as a cause and risk factor[2, 3]. To date, no vaccines or effective therapies are available to treat or prevent this infection[1].

*Cryptosporidium* is a single-celled eukaryote and a member of the phylum Apicomplexa, which also includes the causative agents of malaria and toxoplasmosis. Apicomplexans actively invade cells of their hosts, where many of them replicate within a specialized intracellular compartment formed during invasion called the parasitophorous vacuole (PV)[4]. *Cryptosporidium* occupies a unique intracellular niche in epithelial cells, one that bulges into the lumen of the intestine and is separated from the bulk of the host cytosol[5]. How *Cryptosporidium* embeds itself within the apical cell cortex to maintain its position within the brush border remains poorly understood. Important clues may lie in the ultrastructure of the interface that the parasites build towards the host cell cytosol. This interface is multi-layered and contains a striking array of proteinaceous and membranous structures[6, 7] that likely serve structural as well as physiological roles. However, their molecular composition and mechanism of assembly are largely unknown. Most prominent among these structures is the feeder organelle, a highly convoluted membrane that appears to separate host and parasite cytoplasm. Based on its shape and location, the feeder has long been suspected to represent the site of metabolite exchange between host and parasite. Recent studies on glucose-phosphate transporters have provided molecular confirmation for this hypothesis[8].

*Cryptosporidium* reproduces through asexual and sexual mechanisms, which both lead to the release of motile extracellular forms, merozoites and sporozoites, that will seek out and invade new host cells[9, 10]. Live-cell imaging of sporozoites invading epithelial cells revealed a stereotypical sequence of events, completion of which takes around 5 minutes[11, 12]. Sporozoites glide over the cell surface, until they commit to invasion, which within seconds, leads to polymerization of host actin at the site of contact with the apical end of the parasites[11, 13, 14]. The parasite then envelopes itself with host-cell plasma membrane while rounding up into its intracellular trophozoite form. Electron microscopy studies have captured many aspects of *Cryptosporidium* ultrastructure during invasion. However, the authors differed significantly in their interpretation of these observations, yielding multiple competing models of the invasion mechanism[5, 15, 16]. Following invasion, intracellular parasites further elaborate the host-parasite interface, undergoing growth and schizogony to produce eight merozoites, which egress and continue the cycle[17, 18].

During invasion apicomplexans secrete effectors from specialized organelles[19, 20]. Recently, spatial proteomic studies identified a comprehensive set of putative effectors in *Cryptosporidium* sporozoites[21], including 79 putative dense granule, 24 rhoptry, and 28 microneme effectors. Many of these effectors are delivered to the host-parasite interface[21–24]. The study also identified a previously unrecognized class of secretory organelles unique to *Cryptosporidium,* the small granules. Small granule proteins can be delivered to the interface[21, 24] or exported into the host cell, where they have been reported to facilitate the elongation of microvilli[25–27].

Here, we develop an experimental model to investigate protein secretion and host-parasite interface assembly in living cells. We find that, in *Cryptosporidium*, dense granules are secreted early during invasion, prior to internalization, to establish a cellular holdfast. To our surprise we find that this event uses the conduit formed by the previously discharged rhoptry to access the host cell. Our studies reveal a central and persisting role for the single rhoptry organelle in the establishment of the host-parasite interface by enabling protein translocation and ultimately metabolite exchange. These studies support a unified mechanistic model of *Cryptosporidium* invasion that hinges on effector secretion and the continued remodeling of a remarkably versatile secretory organelle.

## Results

### Poly(ADP)ribose glycohydrolase is a dense granule protein conserved in *Cryptosporidium* and *Toxoplasma*

*Cryptosporidium* secretes more than 150 proteins during and immediately following host cell invasion to secure its intracellular niche[10, 28]. While it is believed that four distinct secretory organelles allow the parasite to deliver effectors with spatial and temporal precision, such secretion has not been directly observed. Most of the secreted effectors studied to date, such as MEDLE2[29] and ROP1[11] show a highly disordered tertiary structure which appears important for their function. Translational fusion of these proteins to structured reporter domains interfered with their trafficking and secretion[29]. To overcome this obstacle, we mined the *C. parvum* secretome for well-folded proteins to facilitate the construction of a real-time secretion reporter. The putative dense granule protein encoded by Cgd8_2160 stood out, as AlphaFold models[30] predicted a highly ordered enzyme fold (Fig. 1A) that showed strong sequence similarity to poly(ADP)ribose glycohydrolases (PARG, interpro domain IPR007724)[21]. This protein has orthologs in *Toxoplasma* and *Eimeria*, but not in *Plasmodium*, *Theileria,* or *Babesia* (Fig 1B). While *Cryptosporidium* has a single PARG-domain containing protein, the *T. gondii* genome encodes two such proteins (Fig. 1B), and spatial proteomic studies using *T. gondii* tachyzoites localized one to the dense granules (280380, TgPARG2) while the other protein (262760, TgPARG1) was found in the nucleus[31].

**Figure 1.**
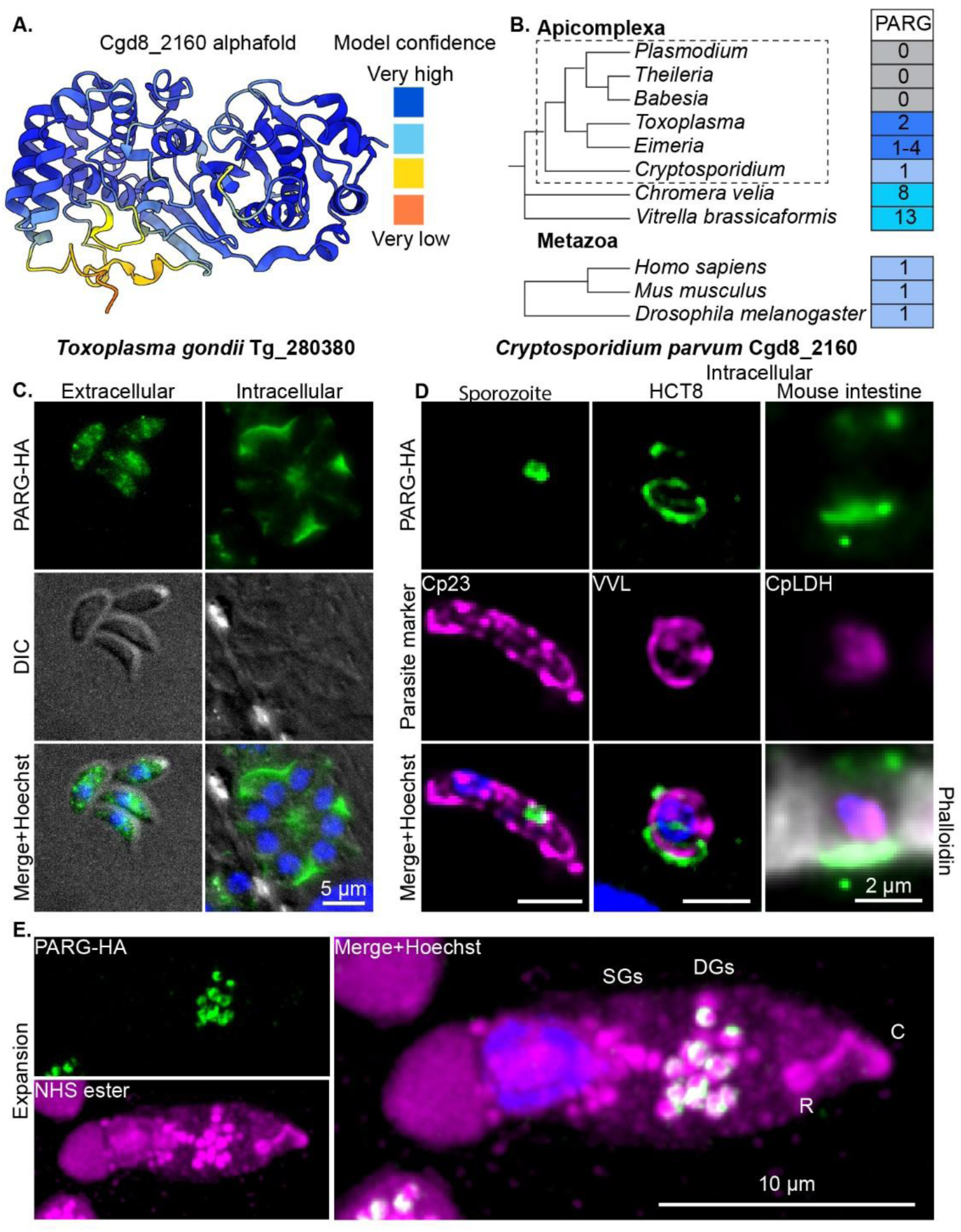
*Cryptosporidium* and *Toxoplasma* express PARG-domain containing dense granule effectors. **A.** Alphafold model of CpPARG colored by model confidence. **B.** Copy number of proteins with predicted poly (ADP-ribose) glycohydrolase (PARG) domains identified by PARG interpro domain annotation search (IPR007724) using VEupath DB. **C.** Immunofluorescence of HA-tagged dense granule PARG (Green) *in T. gondii* (Tg_280380, TgPARG2) in extracellular and intracellular parasites. **D.** Immunofluorescence of *C. parvum* HA-tagged PARG (Cgd8_2160, green) in an extracellular sporozoite and in 1N intracellular parasites, both in *in vitro* in HCT8 cells (24h post-infection) and *in vivo* (9d post-infection). Counterstains are in Magenta. Intestinal sections were also stained with Phalloidin, in gray. **E.** Ultrastructure expansion microscopy (4.5X expanded) showing PARG-HA (green) localization to a population of dense granules labelled with NHS ester (Magenta) in a sporozoite. Small granules (SGs), dense granules (DGs), the rhoptry (R) and the conoid complex (C) are labelled.

We tagged TgPARG2 and CpPARG (PARG) in the respective parasites, inserting a HA-epitope cassette at their endogenous loci using CRISPR-Cas9 (Fig. S1). Immunofluorescence of extracellular *T. gondii* tachyzoites showed numerous granules throughout the parasite cytoplasm, and in intracellular stages TgPARG2 was found secreted into the lumen of the parasitophorous vacuole, labeling the space between the parasites (Fig. 1C).

In *C. parvum* sporozoites, we observed a central patch of staining anterior to the nucleus, a pattern similar to that observed for other dense granule effectors[21] (Fig. 1D). We confirmed this assignment by ultrastructure expansion microscopy of sporozoites, which resolved individual dense granules and labeled the apical portion of NH-ester positive vesicles (Fig.1E, the posterior fraction represents small granules[21]). We performed immunofluorescence on intracellular *C. parvum* in HCT-8 tissue cultures and in sections of the ileum of infected mice. PARG was localized to the host-parasite interface underlying all intracellular stages of *C. parvum* (Figs. 1D, S2). In asexually replicating parasites, new granules containing PARG were abundant in 4N and 8N parasites, consistent with previous studies on dense granule biogenesis[21] (Fig. S2).

To determine whether PARG is required for parasite fitness, we disrupted the gene using CRISPR-Cas9 in *T. gondii* and in *C. parvum*. We obtained both mutants and found no obvious defects when measuring the growth of TgPARG2-KO in culture by plaque assay (Fig. S3A). *C. parvum* PARG-KO parasites showed normal oocysts shedding in infected *Ifnγ^-/-^* mice (Fig. S3B). We conclude that, while the expression of a secretory PARG-domain containing protein is conserved in both parasites, the protein is not required for growth under the conditions tested.

### Dense granule trafficking and secretion during parasite invasion

To observe the behavior of dense granules in living cells, we next engineered reporter parasites that express the green fluorescent protein mNeon fused to the C-terminus of PARG and red fluorescent tdTomato in the cytoplasm. Sporozoites showed the typical central dense granule pattern in green against a red cell body when observed by fluorescence microscopy (Fig. 2A). Infected cells displayed a green interface underlying a red parasite, demonstrating that PARG delivery was not disrupted by the fluorescent protein tag (Fig. 2A).

**Figure 2.**
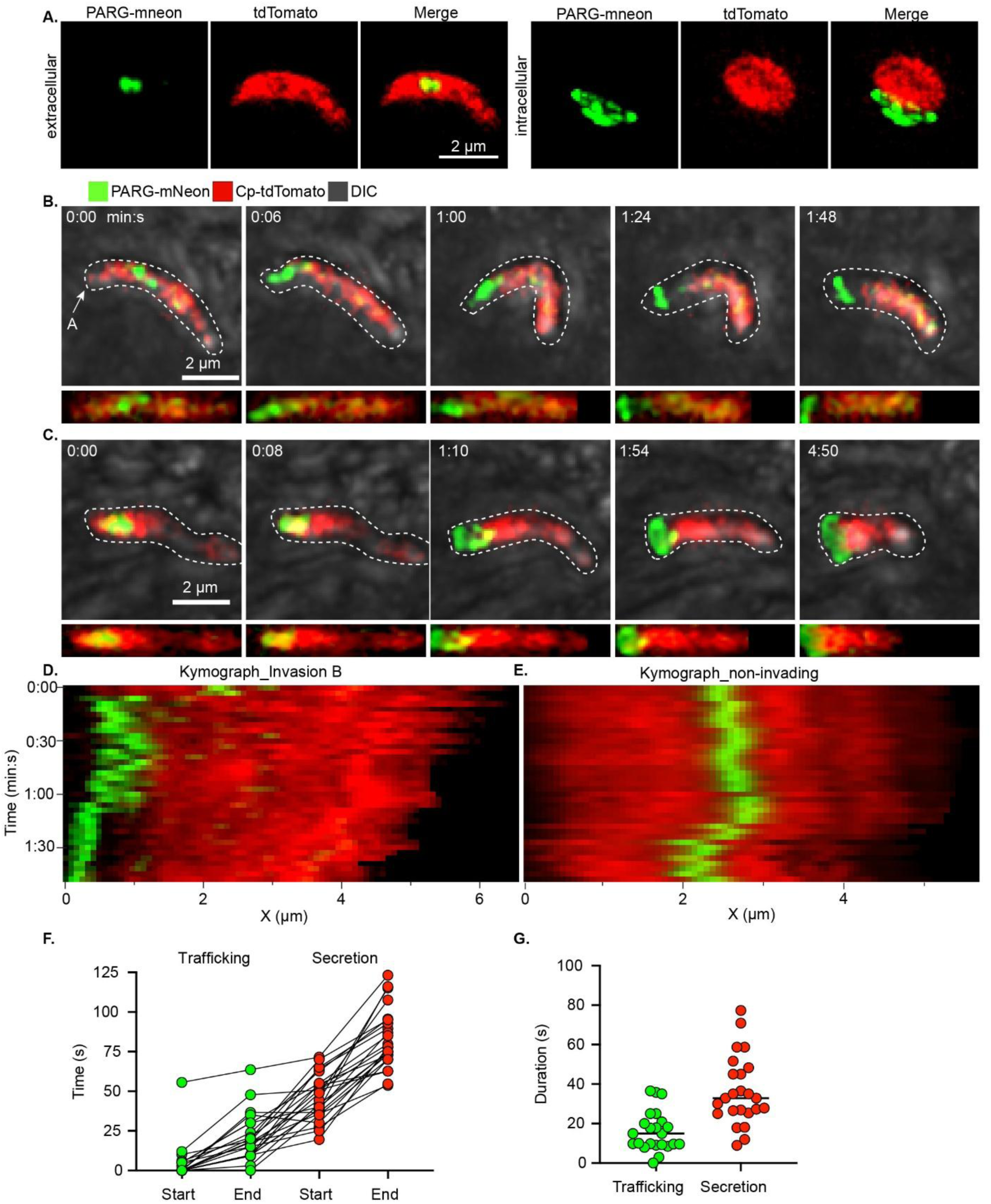
Dense granules undergo coordinated trafficking and secretion during invasion. A reporter strain for live imaging of dense granule secretion. PARG was C-terminally tagged with mNeon, and tdTomato was expressed cytosolically. **A.** Examples of extracellular and intracellular parasites. Reporter sporozoites were used for time-lapse microscopy to visualize invasion. **B-C.** Snapshots of key events in the invasion process from two representative time series. The point of initial contact of the sporozoite with the HCT-8 cell is shown as time 0. Micrographs straightened along the length of the parasite shown below. **D.** A Kymograph from the series in panel B showing the projected fluorescence intensity across the length of the parasite (X-axis) over time (Y-axis). **E.** A kymograph from a movie of a non-invading sporozoite. **F-G.** The timing of the indicated events (N=27). The point of initial contact of the sporozoite with the HCT-8 cell is shown as time 0. **F**. Cumulative timing of selected events relative to T=0 where each line represents an individual invasion event. **G.** The duration of the trafficking (movement of DGs to the apical end) and secretion (deposition of PARG-mneon into the interface) events. Dense granule trafficking took 16 s (± 10) and secretion took 36 (± 17.5 s).

To capture parasite invasion, HCT-8 cells were seeded into glass-bottom chambered slides, mounted on an inverted OMX fluorescence microscope maintaining temperature at 37 °C and CO_2_ at 5% and exposed to fluorescent *Cryptosporidium*. Multi-channel Z-stacks were recorded every 3 to 10 seconds for 10 to 15 min over numerous fields and cultures, resulting in the capture of 27 invasion events (Fig. 2, Fig. S4, Movie S1).

Dense granule secretion proceeded in three steps that were revealed when the PARG-mNeon signal was plotted along the longitudinal axis of the sporozoite as a kymograph (Fig. 2D). Within seconds of stable parasite attachment to the host cell, dense granules were moved ∼1.5 µm from their typical central position to the apex of the sporozoite at speeds of 100 to 200 nm/s (Fig. 2D, Fig. S4A-D). We did not observe such directional movement in non-invading sporozoites (Fig. 2E, Fig. S4E-G, Movie S2). In the second phase, which lasted 60 s on average (Fig. 2F-G), dense granules clustered at the apical end where they appeared to coalesce (Fig. 2A-D). In the final stage, the sporozoite delivered the fluorescent reporter through an apical conduit into the nascent host-parasite interface (Fig. 2B-C). Secretion was completed within 60 s (Fig. 2F-G), after which very little green-fluorescent protein remained in the body of the sporozoite. Overall, invasion took ∼5 min to complete, after which the parasite began to round up (Fig. 2B-D, Fig. S4A-D).

### Dense granule secretion follows host actin polymerization but precedes internalization

*Cryptosporidium* invasion is accompanied by actin polymerization in HCT-8 cells[11–14]. This is initiated early at the site of attachment and is thought to be linked to rhoptry discharge[11]. However, small granule and dense granule proteins have also been found to interact with host-cell actin binding proteins[25–27]. To understand when dense granule secretion occurs in relation to host actin polymerization, we engineered parasites that express a c-terminal PARG-tdTomato fusion (PARG-tdT) and infected HCT-8 cells expressing GFP-lifeact[11]. Recording parasite invasion using time-lapse microscopy, we found F-actin already assembled at the site of attachment when dense granules were still in the middle of the sporozoite (Fig. 3A, Movie S3). We also engineered parasites in which a secreted rhoptry bulb protein (CpRop4-HA) and dense granule protein (PARG-myc) were simultaneously epitope-tagged (Fig. S5A-B). By expansion microscopy, CpRop4-HA was detected already discharged from the rhoptry while PARG-myc-labelled dense granules were still within the parasite (Fig. S5B). Together, these results show that rhoptry discharge and initial actin polymerization precede dense granule secretion.

**Figure 3.**
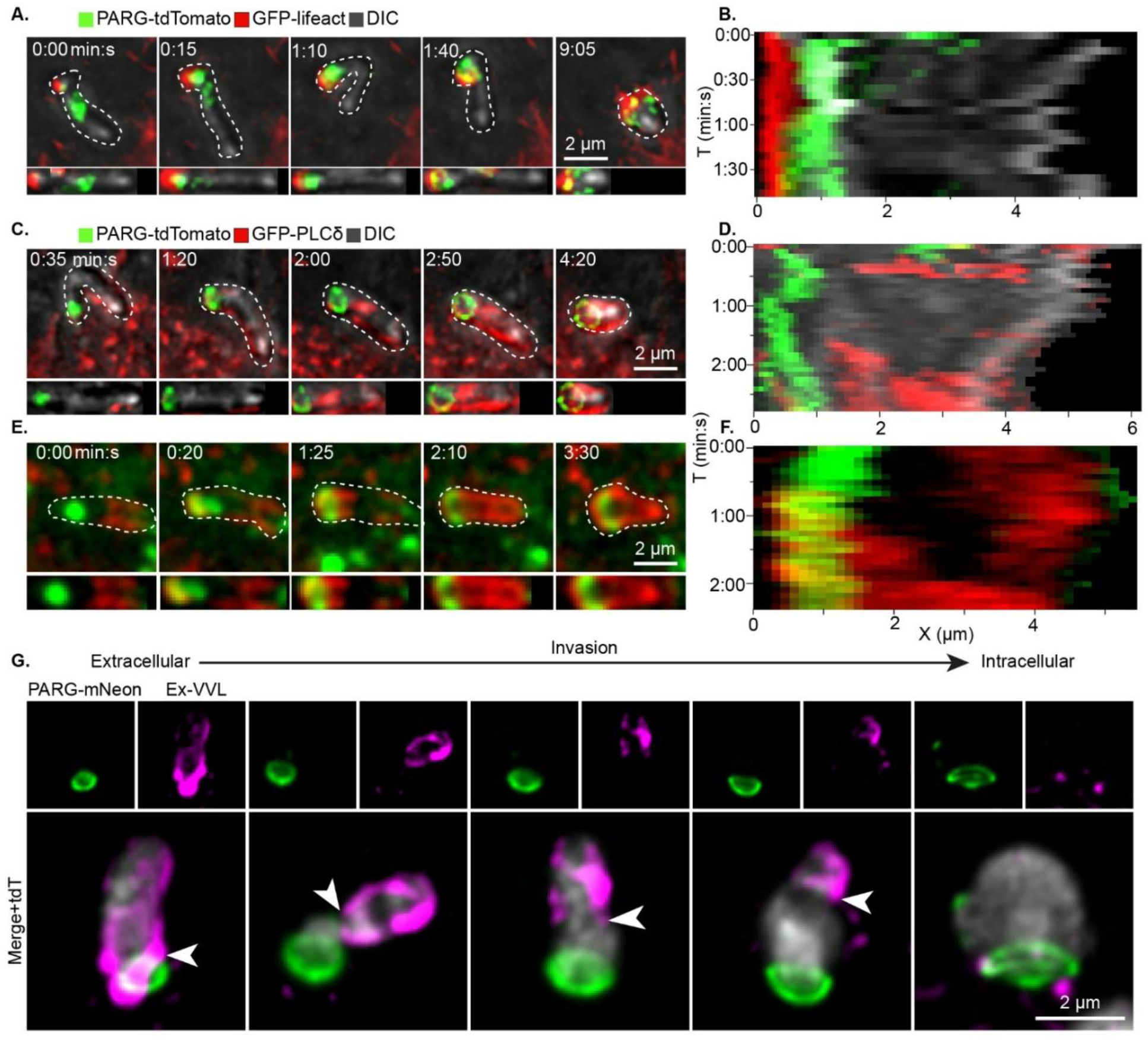
Dense granule secretion occurs after actin polymerization and before internalization. **A-B.** PARG-tdTomato parasites invading HCT8s expressing GFP-lifeact (representative sequence, N=10)**. A.** Snapshots of key scenes. Micrographs were straightened along the length of the parasite and displayed below. **B.** Kymographs showing the projected fluorescence intensity of PARG-tdTomato (green) and GFP-lifeact (red) across the length of the parasite (X-axis) over time (Y-axis). **C-F.** PARG-tdTomato (green) parasites invading HCT8s expressing PLCδ-GFP (red), a plasma membrane marker (representative sequences, N=14). **C-E.** Snapshots of selected scenes highlighting key events. Straightened parasite micrographs are displayed below the original image. **D, F.** Kymographs showing the projected fluorescence intensity of PARG-tdT and PLC-GFP across the length of the parasite for (X-axis) over time (Y-axis). **G.** PARG-mneon:tdT parasites were allowed to invade HCT-8 cells for 2h prior to fixation. PARG-mneon is in green and cytosolic tdT is gray in merged panels. VVL staining (magenta) without permeabilization was used to label the exposed sporozoite surface. Representative images are ordered according to invasion progression. A white arrow marks the boundary between the exposed (VVL-positive) and internalized sections of the parasite.

In *Plasmodium*[32, 33] and *Toxoplasma*[34, 35], dense granule secretion occurs after invasion is completed and is thought to further remodel the host cell and to interfere with cell intrinsic immune responses. We were thus surprised to observe PARG-mNeon secretion within the first minute of host-sporozoite interaction (Fig. 2). To understand this timing relative to parasite internalization we used HCT-8 cells expressing GFP fused to phospholipase C delta (GFP-PLC-δ) to mark the host plasma membrane[11] and invasion of PARG-tdT parasites was recorded by time lapse microscopy. As shown in Fig. 3C-F and Movie S4 the parasite was fully enveloped in host plasma membrane only after the discharge of dense granules. To measure internalization using an orthogonal method, we used a protection assay. HCT8 cells were infected with parasites expressing PARG-mNeon and cytosolic tdTomato (PARG-mn:tdT). Cells were fixed but not permeabilized, and exposed parasite membranes were labelled with biotinylated *Vicia villosa* lectin (VVL). In extracellular sporozoites, this resulted in labelling over the surface (Fig. 3G). In contrast, intracellular parasites were shielded from VVL staining (Fig. 3G). We also observed numerous intermediate stages fixed during invasion, where parasites had secreted PARG and formed the nascent interface, but their surface was still largely exposed (Fig. 3G). We conclude that *Cryptosporidium* invasion proceeds in distinct phases: rhoptry discharge, which triggers F-actin polymerization, followed by dense granule secretion, which generates the initial parasite holdfast within the host cell. Only then, is the parasite engulfed by host membrane resulting in the intracellular trophozoite.

### Dense granules aggregate within the parasite prior to secretion

Next, we used expansion microscopy to understand the structure and fate of dense granule cargo during invasion at high resolution. Oocysts were triggered to excyst, added to HCT-8 monolayers, and fixed 2h later to observe early stages of infection. Expanded cells were incubated with antibodies to detect PARG-HA (green) and amine-reactive NHS-ester conjugated to AlexaFluor-647 to broadly label proteinaceous structures. Sporozoites were observed with their apical end and conoid complex (C) closely associated with host cells (Fig. 4A). When the rhoptry (R) was not yet discharged and visible by NHS ester staining, numerous individual dense granules (DGs) were observed proximal to the parasite nucleus. Following the completion of invasion, parasites were rounded and PARG was deposited at the interface, which was also labeled strongly with NHS ester dye (Fig. 4B).

**Figure 4.**
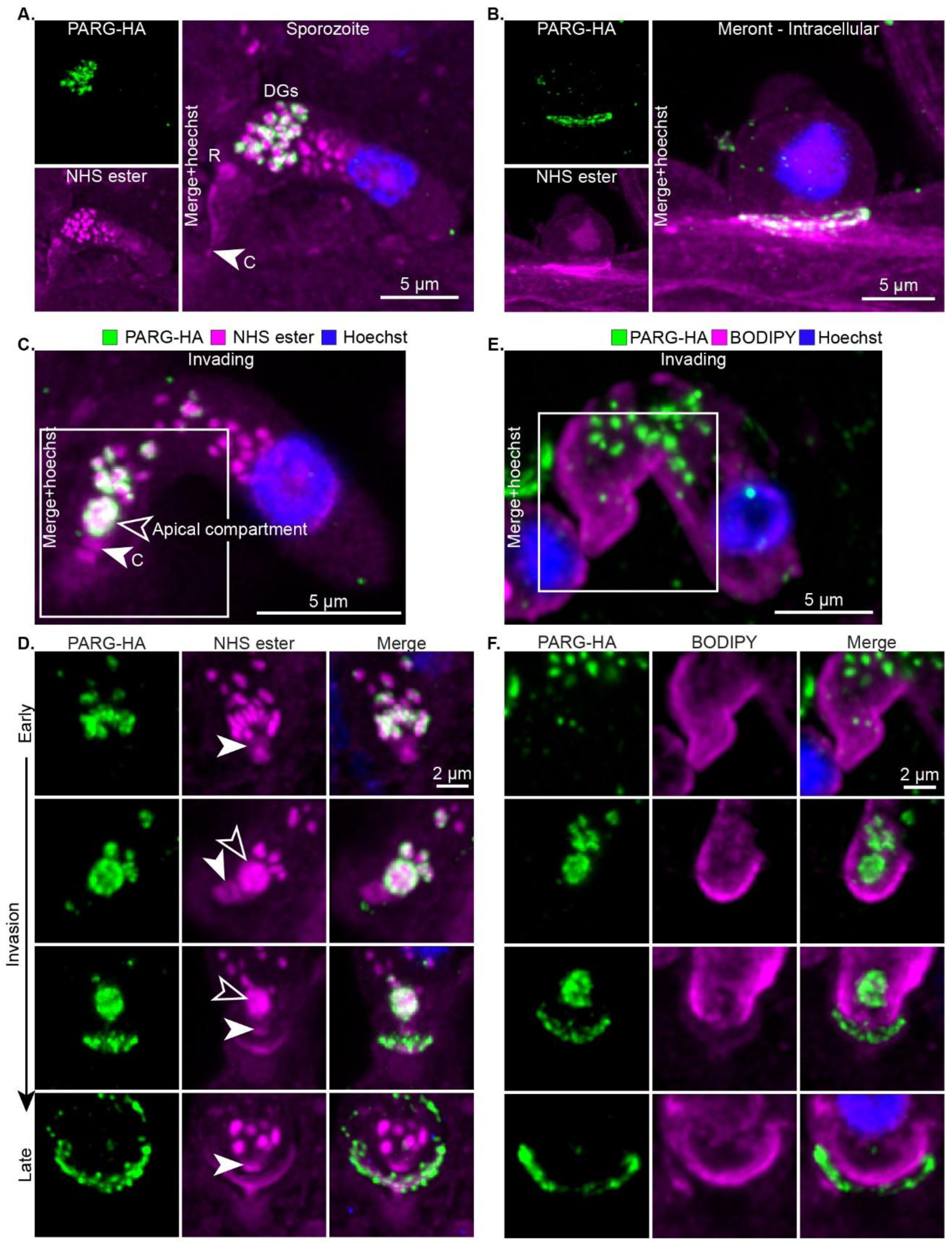
PARG aggregates within an apical compartment prior to secretion into the interface. **A-F.** Ultrastructure expansion microscopy (∼4.5X expansion factor) of invading parasites. NHS ester labelling protein density (**A-D**) or BODIPY labelling membranes (**E-F**) are shown in magenta. PARG-HA staining is shown in green. **A.** Micrograph of a sporozoite interacting with HCT-8 cells. The rhoptry (R) is elongated and connected to the conoid complex (C). The dense granules (DG) are between the nucleus and the rhoptry bulb. **B.** Micrograph of an intracellular 1N meront showing PARG-HA at the interface. **C, E.** Micrograph showing an entire parasite at an early invasion stage. Insets indicate the cropping area for panels **D** and **F**. **C-D.** An open arrow marks the apical compartment, and a filled arrow marks the conoid complex. **D.** Cropped views of the parasite apical end ordered by approximate stage of invasion (early to late) showing the localization of PARG-HA and protein density. **F.** Cropped views of the parasite apical end at different stages of invasion showing the localization of PARG-HA (green) and the parasite plasma membrane stained with BODIPY (magenta). Micrographs showing the entire parasite for panels **D** and **F** are in Fig. S6.

Parasites at different stages of invasion were imaged, and representative examples cropped to highlight the apical end and ordered sequentially according to live imaging data (Fig. 2-3) are shown in Fig. 4C-F and full images and additional examples in Fig. S6A-B. Initially PARG-HA labeled numerous dense granules, but following rhoptry discharge, PARG-HA staining was associated with a single rounded apical compartment with a mean scaled diameter of 0.62 µm (expanded diameter 2.8 µm, Fig. 4C-D, Fig. 6E, S4B). A 3D rendered view of this compartment is shown in Movie S5. The appearance of this structure (open arrow, Fig. 4C-D) coincided with the disappearance of individual dense granules. In subsequent stages, PARG-HA labeled the interface with the host cell (Fig. 4C-D). To define secretion from the sporozoite more rigorously, we also imaged samples counterstained with the membrane dye BODIPY (magenta) (Fig. 4D-F). We note that dense granules coalesced within the boundaries of the sporozoite plasma membrane, and that subsequent secretion from this structure deposited PARG-HA beyond that membrane. Overall, PARG-HA thus moved from the dense granules into an apical compartment, and from there to the interface.

### The apical compartment is membrane-bound with a conduit into the host cell

Some previous ultra-structural studies had described “vacuolation” near the site of attachment in invading parasites[5, 15, 36]. We therefore used electron microscopy to image *C. parvum* invasion in cell culture and infected mice (see methods section for details). We consistently observed an apical electron-translucent compartment in invading sporozoites by transmission electron microscopy (TEM, Fig. 5A-B, S7A-B). This compartment was bound by a membrane (open arrow, Fig. 5A), and electron-dense granules (G) appeared docked to this membrane on the side closer to the parasite nucleus (Fig. 5A). At the apical end (Fig. 5B), we observed a conduit opening into the host and the interface, surrounded by ring-shaped densities (Fig. 5B, S7A-B). Expansion microscopy consistently showed this apical compartment near the conoid complex, visualized using antibodies to tubulin, which labels short sub-pellicular microtubules distal to the conoidal and polar rings[37–39] (filled arrow, Fig. S5C-F).

**Figure 5.**
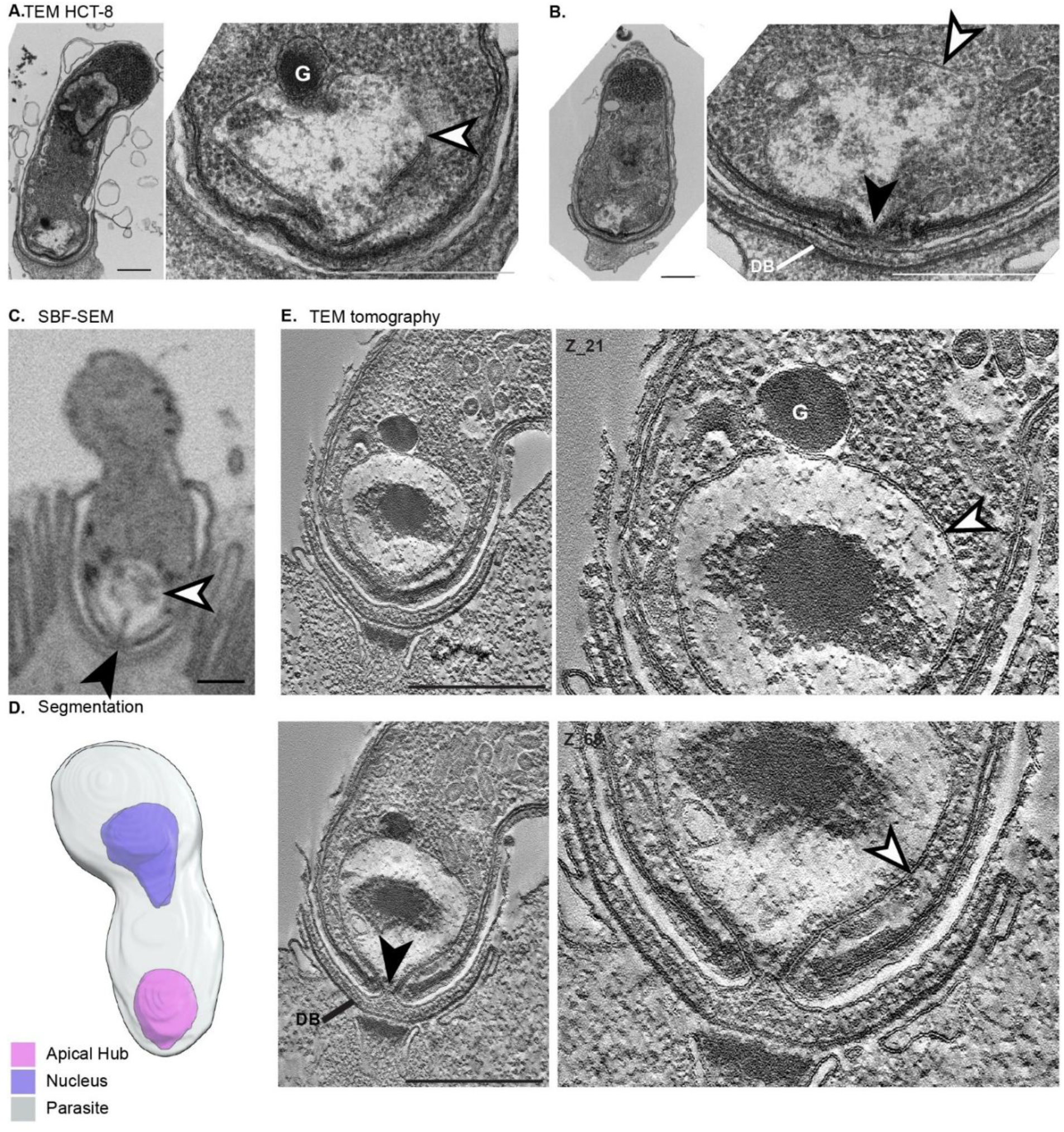
The ultrastructure of the apical compartment. **A-B.** Representative thin-section TEM images of *C. parvum* parasites invading HCT-8 cells. **A.** A lipid bilayer (open arrow) surrounds the apical compartment. An electron-dense vesicle (V) is docked. **B.** The dense band (DB) is formed in the host cell, and a direct conduit (filled arrow) connects the lumen of the apical compartment to the host cytosol. **C-D.** Serial block-face SEM (SBF-SEM) of intestinal sections from infected mice. **C.** A central slice showing an invading parasite. **D.** Segmentation and rendering of the apical compartment (magenta), parasite membrane (gray), and nucleus (blue). **E.** Electron tomography showing an early-stage intracellular parasite invading an HCT-8 cell. The upper panel shows a tomographic section in which an electron-dense granule (G) is docked to the apical compartment. Electron dense material has accumulated inside the compartment. The lower panel shows a different section of the same tomogram revealing a conduit (filled arrow) that connects the lumen of the apical compartment to the host cytosol above the dense band (DB). The dense patch in the host cell below the conduit likely represents the site of initial actin polymerization. Where visible, open arrows mark the boundary or membrane surrounding the apical compartment, and filled arrows mark the conduit connecting the lumen of the apical compartment to the host. Scale bars = 500 nm.

To gain 3-dimensional insight, we turned to serial block face scanning electron microscopy (SBF-SEM)[40] and electron tomography[41]. Movie S6 and Fig. 5C and D show an invading parasite imaged in its entirety by SBF-SEM. This parasite had established the initial interface but was not yet fully internalized. The apical compartment was readily identified and highlighted by segmentation and 3D rendering (Fig. 5D, Fig. S7C, Movie S6). TEM tomography of thicker sections (150 nm) allowed us to follow the membranes at this stage of invasion with greater clarity (Fig. 5E, Movie S7). Importantly, this dataset confirms that the apical compartment is not only close to the apex of the invading parasite, but features a direct conduit from its lumen, through the conoid, and into the host cell where dense band (DB) and the actin patch are readily visible.

### The apical secretory compartment is derived from the rhoptry

Having demonstrated that dense granules merge into an apical compartment prior to secretion, we next considered the origin of this structure. We reasoned that dense granules may fuse with the membrane of the empty rhoptry following its initial discharge. Invasive *Cryptosporidium* stages harbor a single rhoptry that injects its cargo via a secretory apparatus at the apical tip directly into the cytoplasm of the host cell[11, 37]. To test this hypothesis using molecular markers, we identified *Cryptosporidium* with sequence similarity to rhoptry membrane proteins previously characterized in other apicomplexans [11]. TgRON11 is a protein with 7 transmembrane domains that localizes to the membrane of the rhoptry neck in *Toxoplasma*[42] and we identified its homolog, CpRON11 as Cgd3_2010. We also selected a homolog of TgARO, a protein with armadillo repeats that in *Toxoplasma* is tethered to the rhoptry membrane via N-terminal acylation[43] (CpARO, Cgd2_370) and engineered *C. parvum* strains carrying a HA-epitope tag in each of these proteins.

When observing sporozoites using expansion microscopy where the single rhoptry is readily identified by NHS-ester staining[21], we found that labeling for CpARO extended to the entire rhoptry and that CpRON11 was restricted to a region of the rhoptry neck (Fig. 6A-B, standard immunofluorescence in Fig. S9A-B). Next, we used both strains to visualize the fate of the rhoptry membrane during and following invasion (Fig. 6C-D, S8). We found that both proteins decorated the periphery of the apical compartment (Fig. 6C-D, apical views, open arrow in NHS ester panels, S8 full views), suggesting that, prior to secretion, dense granules fuse with the empty rhoptry. To test this directly, we counterstained with a rabbit antiserum to *Cryptosporidium* dense granule protein 8 (DG8)[22]. This protein is localized to dense granules in sporozoites and forms a ring at the interface following invasion[22] (Fig. 6C-D, S8). We observed a compartment bound by a membrane decorated with CpARO and CpRON11, while we detected DG8 in the lumen of this compartment. We quantified the diameter of the CpARO and CpRON11-positive compartments in invading and immature intracellular parasites and found that their scaled maximum (Feret’s) diameter (0.7 µm CpARO, 0.662 µm CpRON11) was comparable to the diameter of the structure visualized by electron microscopy and by NHS ester staining (Fig. 6E). We conclude that the rhoptry not only secretes its own cargo but is retained and remodeled to subsequently coordinate the secretion of dense granules.

**Figure 6.**
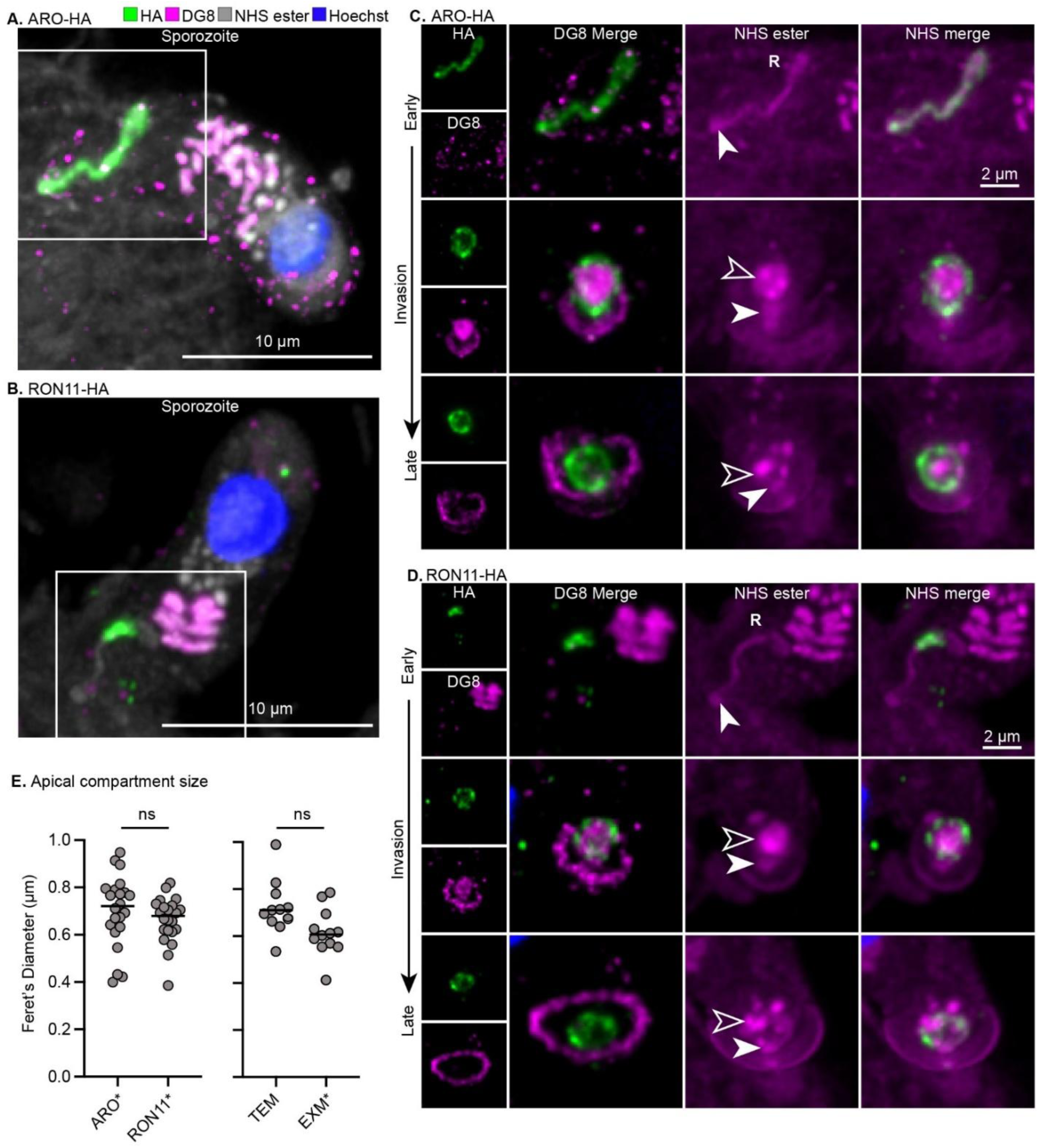
The apical secretory compartment is derived from the rhoptry. **A-D.** Expansion microscopy of invading sporozoites. **A-B.** View of sporozoite interacting with HCT-8 cells. Dense granule cargo is labelled with anti-DG8 sera (magenta), and protein density (NHS ester) is in grayscale, Hoechst is in blue. The rhoptry membrane markers CpARO (**A,C**) and CpRON11 (**B,D**) were epitope-tagged and stained (green). **A.** CpARO labels the entire rhoptry (R). **B.** CpRON11 labels the distal section of the rhoptry neck. The white square indicates the cropping area for figures **C** and **D. C-D.** The apical end of parasites at different stages of invasion. Left-most panels show individual channels for HA and DG8. Larger panels to the right show a DG8-HA merge, NHS ester, and an NHS-HA merge, respectively. Images are arranged according to invasion stage. **C.** CpARO-HA expressing parasites. CpARO labels the periphery of the apical compartment indicated with an open arrow and is connected to the conoid complex marked with a filled arrow in NHS images. **D.** CpRON11-HA expressing parasites. CpRON11 labels the periphery of the apical compartment. Micrographs showing the entire parasites in panels C-D are in Fig. S8. **E.** Quantification of the apical compartment maximum (Feret’s) diameter captured by various imaging modalities and markers. The diameters were not significantly different (p > 0.05, unpaired Welch’s T-test). Asterisks indicate where the diameter was scaled based on an estimated expansion factor of 4.5. The mean diameters are as follows: EXM_scaled (NHS ester) = 0.62 ± 0.100 µm (2.77 µm expanded), TEM = 0.72 ± 0.12 µm, CpARO_scaled = 0.70 ± 0.15 µm (3.15 µm expanded), CpRON11_scaled = 0.66 ± 0.10 µm (2.98 µm expanded).

### The rhoptry membrane is the precursor of the feeder organelle

We wondered whether dense granule secretion concludes the non-canonical use of the *Cryptosporidium* rhoptry, or whether this compartment might serve additional functions as the parasite builds its intracellular niche. To explore this idea, we stained intracellular stages 2 and 6 hours post-infection for the rhoptry membrane makers CpARO and CpRON11. Labeling of CpRON11 and CpARO persisted and morphed into a linear structure at the interface of parasite and host (Fig. 7A-D, S10 expansion, S9 standard immunofluorescence). In more mature trophozoites, this region of CpRON11 and CpARO staining was found to coincide with the membrane dye BODIPY that separated host from parasite at the interface and localized just above the dense band (filled arrow) labeled by NHS-ester (Fig. 7C-D). This CpARO-HA positive structure lengthened between 2 and 6h post-infection, and this growth had a linear relationship with the growth of the nucleus (the nucleus was previously shown to grow as parasites develop[17], Fig. S7C).

**Figure 7.**
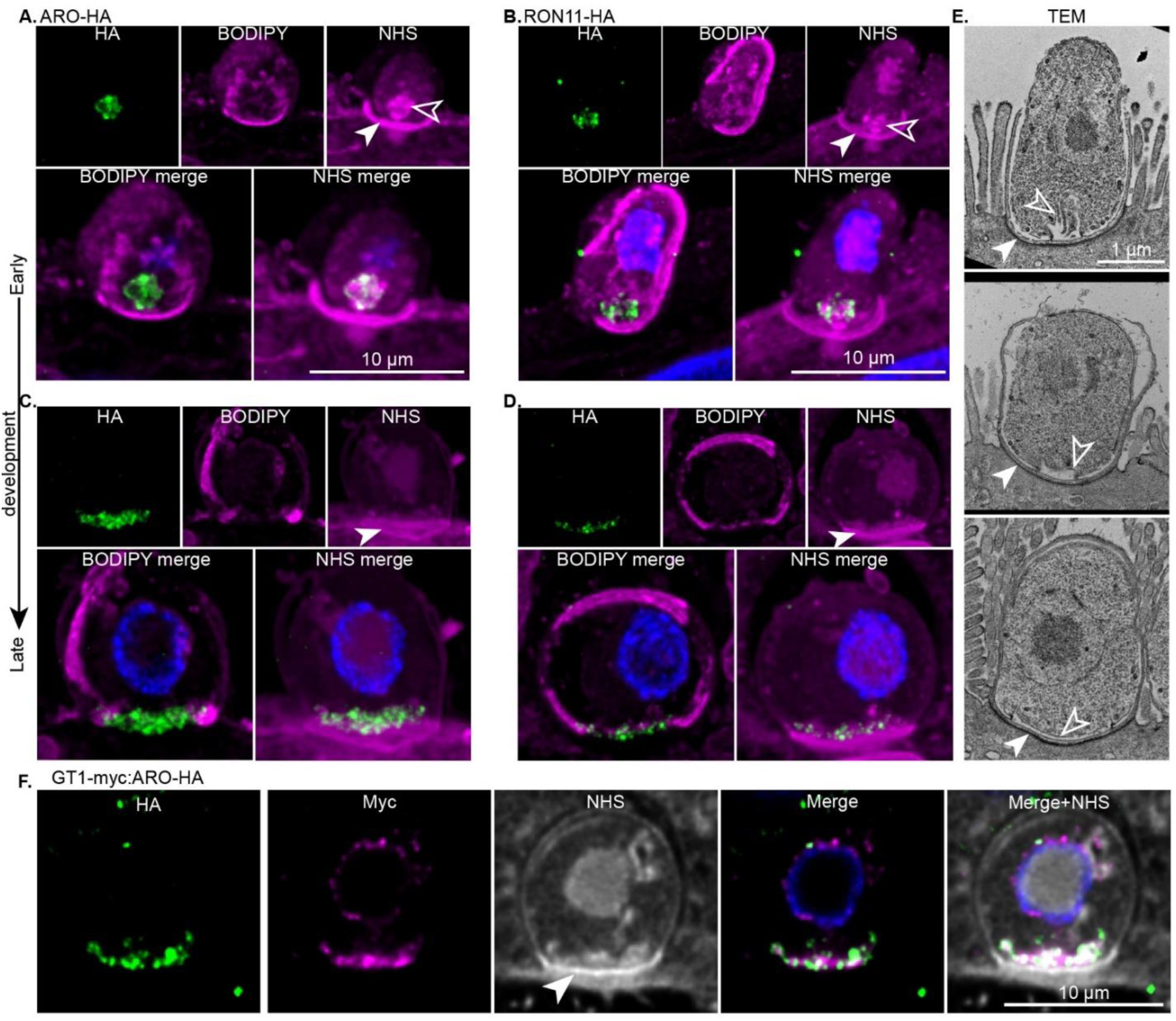
The rhoptry membrane develops into the feeder organelle. **A-D.** Expansion microscopy of intracellular parasites expressing CpARO-HA (**A, C**) or CpRON11-HA (**B, D**) in HCT-8 cells. Upper panels show individual channels (HA in green, BODIPY and NHS ester in magenta). Lower panels show BODIPY and NHS ester channels merged with the HA channel and Hoechst (blue). **A-B.** Immature intracellular parasites that are still slightly elongated and smaller. The apical compartment or feeder organelle is marked by an open arrow, and the dense band is marked by a filled arrow in NHS images where visible. **C-D.** Mature intracellular parasites that are more rounded and larger. The apical compartment is no longer visible. CpARO (**C**) and CpRON11 (**D**) both localize to the interface above the dense band (filled arrow) visible by NHS ester staining, and between the two arms of the parasite plasma membrane shown by BODIPY. **E.** Examples of parasites at different stages of feeder organelle development from thin section TEM of infected mouse tissue. Open arrow = apical compartment/feeder membrane, filled arrow = dense band. **F.** Expansion microscopy of intracellular parasites co-expressing GT1-myc and ARO-HA. GT1 and ARO co-localize at the interface above the dense band. Some perinuclear GT1 is visible, likely due to transit through the secretory pathway. Panel 4 shows a merge of GT1-myc, ARO-HA and Hoechst. NHS ester (gray) is shown in the right-most panel.

We sought additional ultrastructural insight into this transition. Intestinal sections from *C. parvum* infected mice were analyzed using thin-section TEM and SBF-SEM. Representative TEM and SBF-SEM images are shown in Fig. 7E and S10 and full SBF-SEM volumes and rendered segmentation in Movie S8. The structure that matches the persisting rhoptry membrane is the membrane of the feeder organelle (7E, S10 open arrow). This membrane separates host and parasite, is found just above the dense band (filled arrow), and grows laterally along with the intracellular development of the parasite[5, 6, 8]. Like the membrane of the former rhoptry, the feeder membrane emerges from a ring structure linking the plasma membranes of both host and parasite (Fig. 7E, S10).

To rigorously test the hypothesis that the rhoptry membrane transitions into the feeder membrane, we engineered a new parasite strain to simultaneously tag both structures with a molecular marker. In a recent study, Xu and colleagues elegantly demonstrated that the glucose transporter CpGT1 is localized to the feeder organelle in intracellular parasites[8]. We thus engineered parasites in which GT1 was c-terminally tagged with the c-myc epitope by homologous recombination at its endogenous locus. This insertion carried a second epitope cassette encoding CpARO-HA under its own promoter (GT1-myc:ARO-HA, Fig. S1). We detected both proteins simultaneously by immunofluorescence (Fig. S9D) and expansion microscopy (Fig. 7F) and found that they label the same structure. In mature intracellular parasites, both proteins co-localized at the host parasite interface above the dense band labelled by NHS ester (Fig. 7F). We conclude that the membrane of the rhoptry persists as the initial boundary between parasite and host and through subsequent remodeling gives rise to the feeder organelle membrane.

## Discussion

Apicomplexa are masters of cell invasion and intracellular parasitism. They not only penetrate host cells but remodel them extensively to hijack host organelles, enable metabolite uptake, and broadly disable mechanisms of cell intrinsic defense and immunity[4]. This is accomplished by an army of effector proteins delivered by multiple secretory organelles[19] most extensively studied in in *Toxoplasma* and *Plasmodium*. Both these parasites establish a vacuole deep within the host cytosol[19]. In contrast, *Cryptosporidium* is restricted to the apical surface of epithelial cells[9, 10]. These highly polarized cells harbor specialized structures, including tight junctions and microvilli[44, 45]. Recent studies have found that several of these features are subject to manipulation by a rapidly growing list of *C. parvum* effectors[44–46].

By observing dense granule behavior in real-time during *Cryptosporidium* sporozoite invasion, we found coordinated and directed trafficking toward the site of attachment, at a rate consistent with active transport along cytoskeleton filaments[47] (Fig. 2). In intracellular *T. gondii* parasites, dense granules are anchored to actin filaments, and their transport requires Rab11A[48] and MyoF[49]. Further investigation is needed to determine the mechanism of dense granule trafficking in *Cryptosporidium*, in particular how relocation is triggered as part of the invasion process.

The machinery associated with rhoptry and microneme secretion appears conserved across the Apicomplexa[20]. Discharge proceeds through the conoid, an apical cone comprised of unique tubulin polymers[50]. Micronemes secrete adhesins in a calcium-stimulated process, and when displayed on the extracellular surface, these proteins enable attachment and actin/myosin-based gliding motility[51],[52]. Next, contact of specific microneme proteins with their host cell receptors triggers secretion of rhoptries[53], which deliver their contents into the cytoplasm and the plasma membrane of their target cell[54]. The discharge of rhoptry contents relies on a complex molecular machine, including the rhoptry secretion apparatus, allowing rhoptry proteins to traverse three membranes, the rhoptry membrane and the plasma membranes of parasite and host[37, 55].

The mechanisms by which dense granules are secreted appear less conserved. In *Toxoplasma,* dense granule exocytosis occurs through specialized annular rings (apical annuli) that create gaps in the inner membrane complex to allow fusion with the plasma membrane[56, 57]. These annuli are also found in *Plasmodium*[58]. However, electron microscopy failed to detect equivalent structures in *Cryptosporidium* sporozoites [5, 15, 59]. While the *C. parvum* genome encodes orthologs of some *Toxoplasma* proteins found in annuli (e.g. LMDB3), numerous critical components, including AAP1 to 7, are missing. Our studies suggest a distinct mechanism of dense granule discharge in *Cryptosporidium* where dense granules (and potentially small granules) dock to and fuse with the membrane of the empty rhoptry (Fig. 8A). The molecular mechanisms regulating recognition of and fusion with the rhoptry membrane are still unknown. Ion fluxes triggered by host contact and rhoptry exocytosis were described to regulate *T. gondii* dense granule secretion[60]. Similar processes may activate motor proteins, GTPases, and fusion machinery in *Cryptosporidium*. Once proteins cross the rhoptry membrane, the pore established during rhoptry discharge may bypass the need for additional secretory structures. A conduit connecting the lumen of the apical compartment and the host cytosol is clearly visible in many of our images (Fig. 5). Whether this channel is formed by persisting elements of the rhoptry secretion apparatus or is formed from other components during invasion remains to be explored.

**Figure 8.**
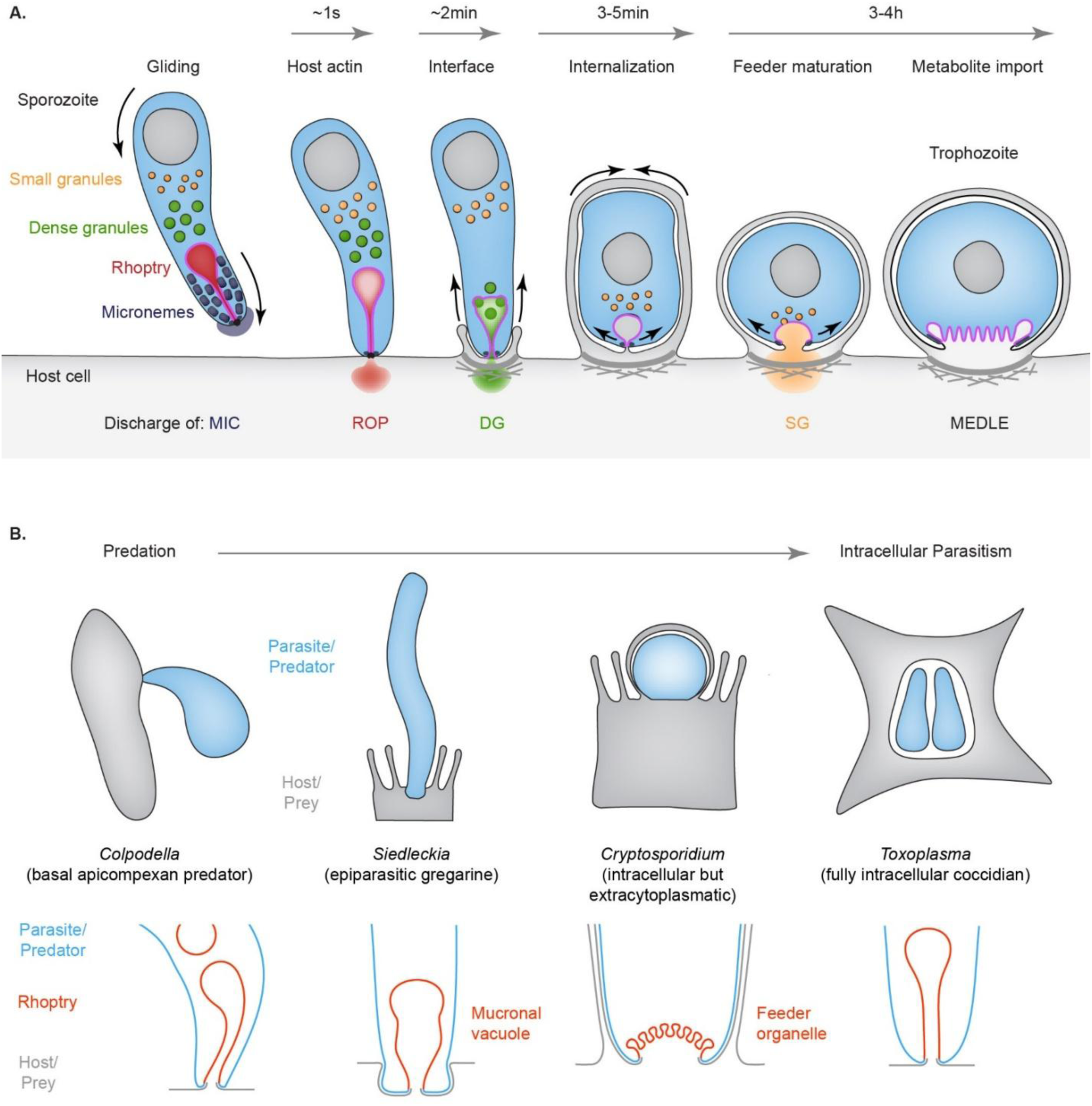
Updated model for Cryptosporidium invasion and effector secretion. **A.** We propose the following model for Cryptosporidium invasion and effector secretion and establish the timing of selected events above the cartoon. First, the parasite establishes stable contact with the host cell and inserts its apical end into an invagination in the host cell membrane. Then, rhoptry discharge and host actin accumulation at the site of attachment occurs rapidly, within 5 seconds of initial contact. The dense granules then migrate to the apical end, a process which takes about 10 seconds. They fuse at the apical end and are secreted into the host-parasite interface. Once the interface has been established, the parasite engages the host plasma membrane and forms its parasitophorous vacuole. The parasite further rounds up and matures. The rhoptry membrane then develops into the feeder organelle starting at three to four hours after invasion. **B.** Model of the evolution of intracellular parasitism in Apicomplexa. *Colpodella* is a predator of other protists, feeding by cellular vampirism or myzocytosis. *Siedeleckia* is a gregarine parasite of marine worms that inserts an apical holdfast and feeding structure called mucron into its host cell but remains largely extracellular, and *Toxoplasma* is a fully intracellular parasite growing within a parasitophorous vacuole in the cytoplasm of the host cell. All these organisms feature a rhoptry (red). In *Toxoplasma* this is solely an invasion organelle, but it appears to have evolved from an organelle that combined host penetration with nutrient acquisition.

Multiple dense granule proteins are translocated into the host cell by *Plasmodium* and *Toxoplasma* in a two-step process[20, 61]. First, dense granule proteins are delivered into the PV by exocytosis. These proteins then cross the PVM through a dedicated translocon[62]. Several reports have demonstrated equivalent export of *Cryptosporidium* dense granule[27], small granule[25, 26], and other[29] proteins into the host cell. Translocated proteins are typically processed by aspartyl proteases, and disruption of this processing leads to their entrapment in the parasite[25, 27, 29]. Only one component of the previously described *Plasmodium* and *Toxoplasma* translocons (ClpB2/HSP101), is found encoded in the *Cryptosporidium* genome, but its function remains unknown[21]. Our model predicts the rhoptry/feeder membrane to be the only membrane separating parasite and host at the interface. Fusion of granules with this membrane thus delivers their contents topologically into the host cytosol. However, many secreted proteins remain restricted to the interface[21, 24, 63] (Fig. 4). This may be because they engage other host and parasite proteins to form higher-order structures that retain them. Alternatively, the two proteinaceous plates, the dense band and the filamentous mesh (Fig. S10E) [6, 10], that underly this membrane may establish a barrier. This barrier may be permeable to metabolites, allowing the parasite to feed, but impede the movement of proteins, potentially shielding the parasite and its membrane from certain types of immune-mediated restriction[64]. Further research is needed to understand the structure, composition and physiology of these non-membranous components of the interface.

Dense granules in *T. gondii*[34, 35, 65] and *Plasmodium spp*[33, 66] are discharged only after penetration of the host cell has concluded and the parasite is surrounded by the vacuole. Surprisingly, we found that *Cryptosporidium* secretes the contents of its dense granules prior to internalization (Fig. 3C-F). *Cryptosporidium* builds an intricate host-parasite interface which has been described by electron microscopy[6, 15] to contain multiple layers of protein plates anchored to membranes (Fig. S10E). Recent work by multiple laboratories has highlighted proteins secreted from rhoptries, small and dense granules as likely components of these structures[11, 21–24, 63]. The findings of this study suggest that *Cryptosporidium* uses the secretion of rhoptry and dense granules to first establish a holdfast in the host cell, which includes the dense band. Only then, in a second step is the parasite engulfed by host membrane (Fig. 3). We believe that this reflects the gradual evolution of intracellular parasitism in apicomplexans (Fig. 8B). As has been suggested by others as well[19, 67], these protists may have adapted organelles initially used for predation to parasitism. Rhoptries are shared cellular features of predators like *Colpodella,* which uses cellular vampirism or myzocytosis to aspirate the cytoplasm of prey cells[68] as well as gregarines like *Siedleckia*[69], *Selendium*[70] or *Gregarina*[15], which parasitize the epithelia of invertebrates while remaining largely extracellular. Gregarines establish an intracellular holdfast called the mucron, akin to the haustorium of mistletoe (Fig. 8B). Structural similarities between gregarines and *Cryptosporidium* have been previously noted, as has their close phylogenetic relationship[15, 71, 72]. We suggest that the rhoptry not only represents a shared secretory organelle, but through persistence and transformation gives rise to the specialized interface structures found in apicomplexan parasites.

In *Cryptosporidium*, we provide molecular evidence that the rhoptry and its membrane persists well beyond invasion. This organelle is remarkably malleable in its function, first injecting its own cargo to initiate invasion, then serving as a hub and conduit for the secretion of additional secretory organelles and finally serving as the membrane of the feeder organelle. The metabolic capabilities of *Cryptosporidium* are stripped down to a bare minimum, and the parasite relies heavily on import of host metabolites[9, 10, 73]. Multiple recent studies have highlighted a wide range of import mechanisms[8, 74, 75]. When compared to *Toxoplasma* or *Plasmodium*, *Cryptosporidium* shows a significant expansion of its repertoire of transporters[76], one of which was recently been localized to the feeder membrane[8].

*Cryptosporidium* host cell invasion has been puzzling with an abundance of complex structures and multiple competing models as to how they are formed. A particularly vexing question has been how the process may end up with a single membrane separating host and parasite, the feeder. Our findings support a unifying and satisfyingly simple mechanistic model outlined in Fig. 8A. The rhoptry and its continuous transformation are central to this model. Invasion of all Apicomplexa requires an initial pore in the host cell membrane. This pore is generated by elements of the highly conserved rhoptry secretory apparatus, in a way that remains poorly understood[20, 54]. One challenge in better understanding this apparatus, even in facile model systems like *Toxoplasma*, is the transient nature of its deployment during rhoptry secretion and invasion, which occur rapidly in *Toxoplasma* and *Plasmodium*. Our model implies that aspects of the rhoptry and potentially the early pore and/or host parasite junction persist past invasion, and this persistence may make them more tractable. This structure is notable as a protein rich ring joining the rhoptry membrane with the plasma membranes of both host and parasite. Understanding the topology, composition and genesis of this ring is not only central to understanding *Cryptosporidium* invasion but may also hold important clues for the mechanism of host cell invasion across Apicomplexa.

## Acknowledgements

We are grateful to Dominique Soldati and Sebastian Lourido for the kind gift of plasmids for genetic manipulation of *T. gondii*, and thank Jessica Byerly, Chloe Tang, Gracyn Buenconsuejo, Abby Daniels, and Aurelia Balestra for materials and help with animal experiments. This work was supported in part by grants from the National Institutes of Health to B.S. (R37-AI112427 and R01AI127798), to A.C. (F32AI183654), and to the Penn Vet Imaging core (S10 OD032305) and by support from the Robert Koch-Institute to T.B., M.L., and C.K.

## Author contributions

A.C. and B.S. conceived the study. A.C. generated transgenic parasites with contributions from A.G. A.C. performed and analyzed light microscopy experiments and prepared fixed infected cells for electron microscopy. T.B. performed additional staining and embedding of both mouse tissue and infected cells and performed electron microscopy experiments with support from M.L. and C.K. T.B. and A.C. performed segmentation of SBF-SEM volumes. A.C and B.S. wrote the manuscript with contributions from all authors.

## Supplementary figures and legends

**Figure S1.**
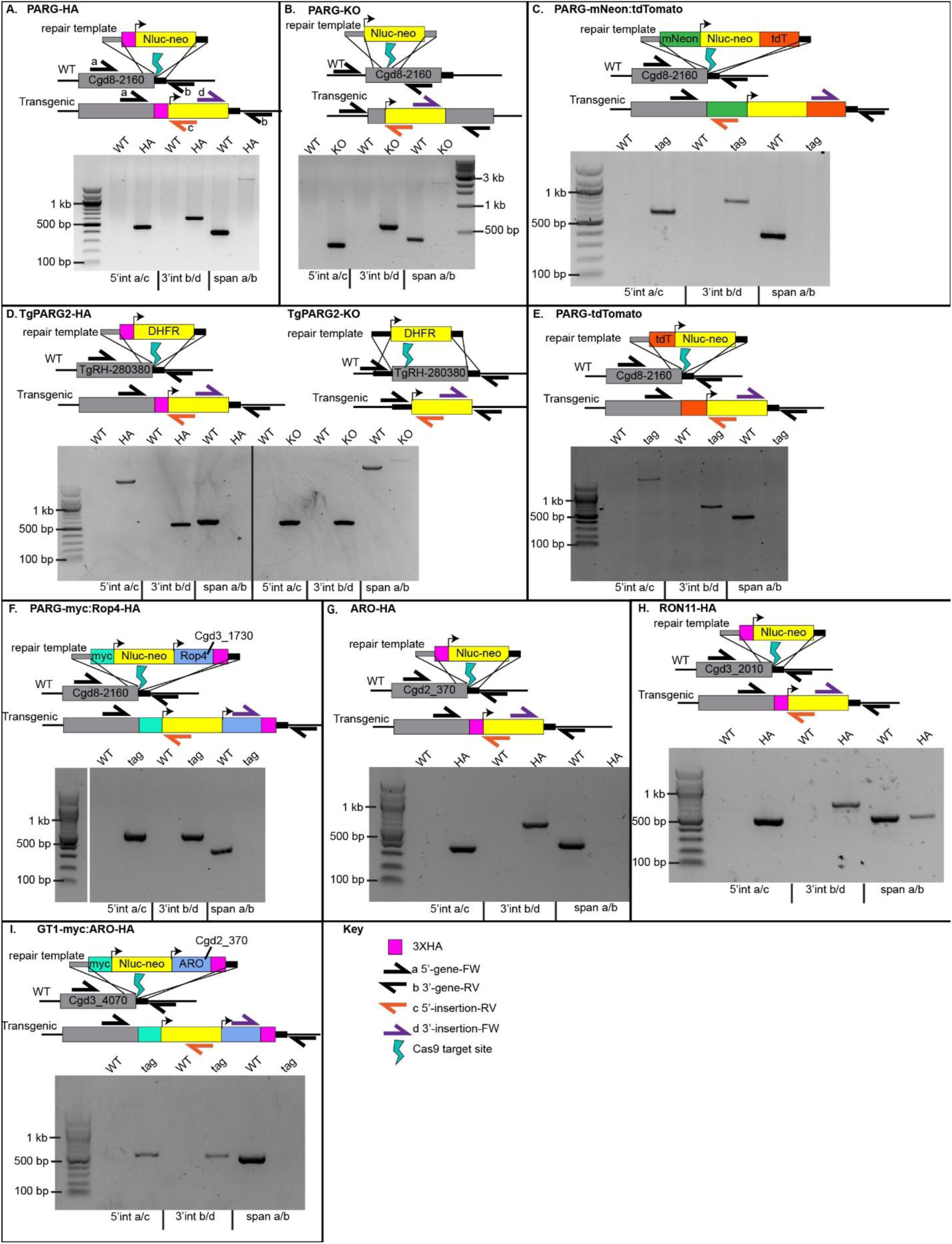
Strain generation. CRISR-CAS9 targeting and homology-directed repair design are shown in a schematic. Promoters are annotated as bent arrows, and primers for checking integration of the construct are annotated as split arrows. See Table S3 for more details on the primers used for integration PCR and the expected amplicon sizes for each transgenic. The gel corresponding to integration PCR for each transgenic is shown below the schematic in inverted contrast. The sample (WT or transgenic at the indicated locus) are labelled at the top of the lane, and the PCR/primers used are labelled at the bottom. Key ladder markers are annotated on the left or right of the image. Transgenic lines are as follows: **A.** PARG (Cgd8_2160) HA epitope tagging **B.** PARG-KO – nluc-neo is knocked into the 5’ end of the ORF, excising half of the PARG domain and splitting the ORF **C.** PARG-mneon:tdTomato – PARG tagging with mNeon and cytosolic expression of tdTomato attached to nluc-neo via a self-cleaving T2A peptide. **D.** *T. gondii* dense granule PARG (280380, TgPARG2) HA epitope tagging and TgPARG2 knockout by gene replacement **E.** PARG-tdT – PARG tagging with tdTomato **F.** PARG-myc:ROP4-HA – PARG is myc-tagged at its endogenous locus and CpROP4-HA is knocked in downstream of the nluc:neo cassette with 500bp of sequence upstream of the start codon to include the native promoter **G.** ARO (Cgd2_370) HA epitope tagging **H.** RON11 (Cgd3_2010) epitope tagging. **I.** GT1 (Cgd3_4070) myc-tagging with ARO-HA (Cgd2_370) with 500bp of sequence upstream of the start codon is knocked in downstream of the nluc-neo cassette.

**Figure S2.**
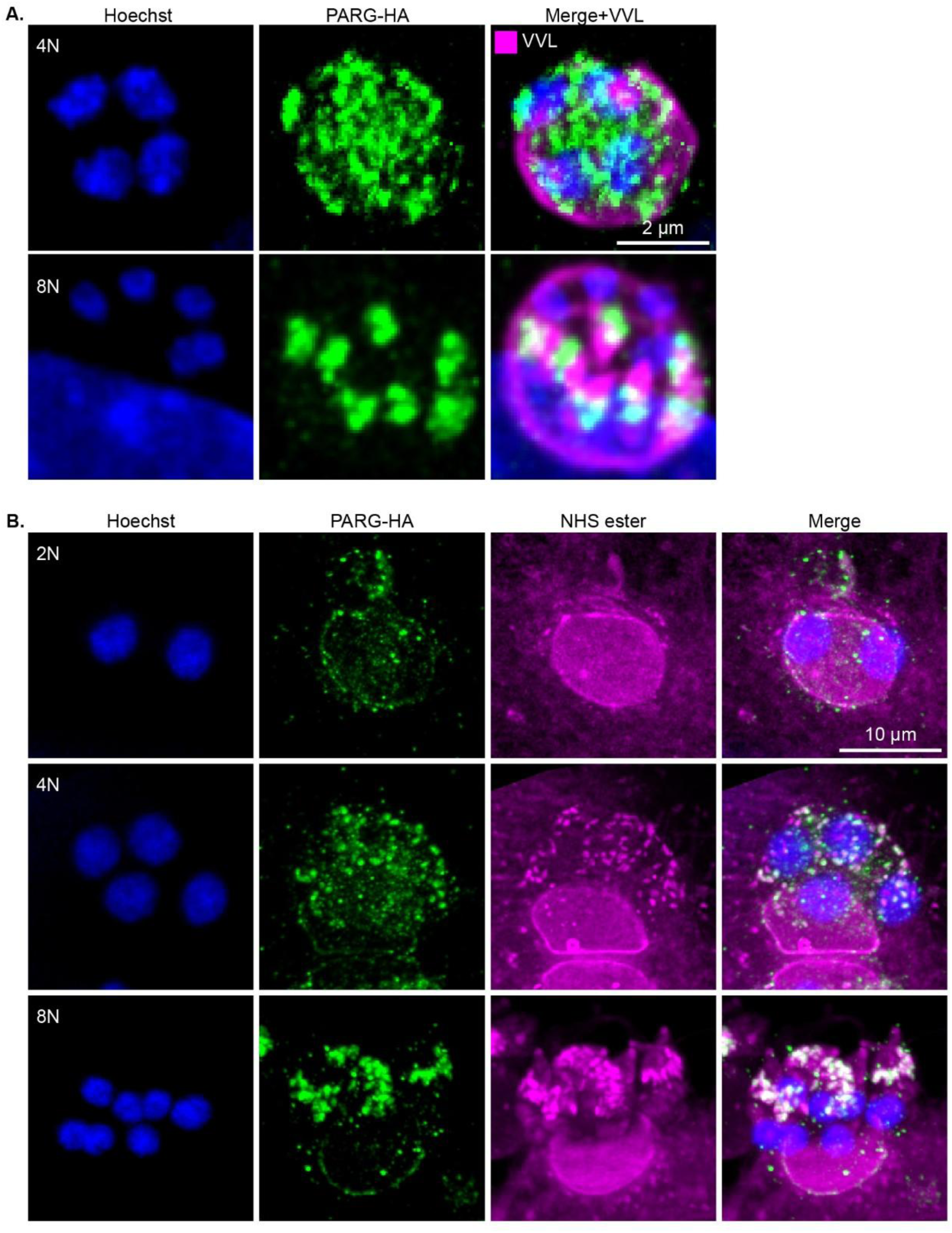
Dense granule synthesis occurs after 2 nuclear divisions. **A.** Immunofluorescence microscopy showing the distribution of PARG-HA in parasites undergoing asexual replication in HCT-8 cells. Newly synthesized PARG is detected in intracellular parasites starting after 2 nuclear divisions (4N). At the third division, when the parasites have 8 nuclei, the PARG-HA signal coalesces into discrete puncta located anterior to the nucleus of the budding merozoites. **B.** Expansion microscopy of parasites undergoing asexual replication showing PARG-HA staining, protein density, and number of nuclei in 2N, 4N, and 8N meronts. Increased PARG-HA expression was observed in 4N meronts. The individual PARG-HA-positive puncta were smaller and more numerous in 4N than in 8N parasites. The intensity of PARG-HA staining at the dense band did not increase noticeably during merogony.

**Figure S3.**
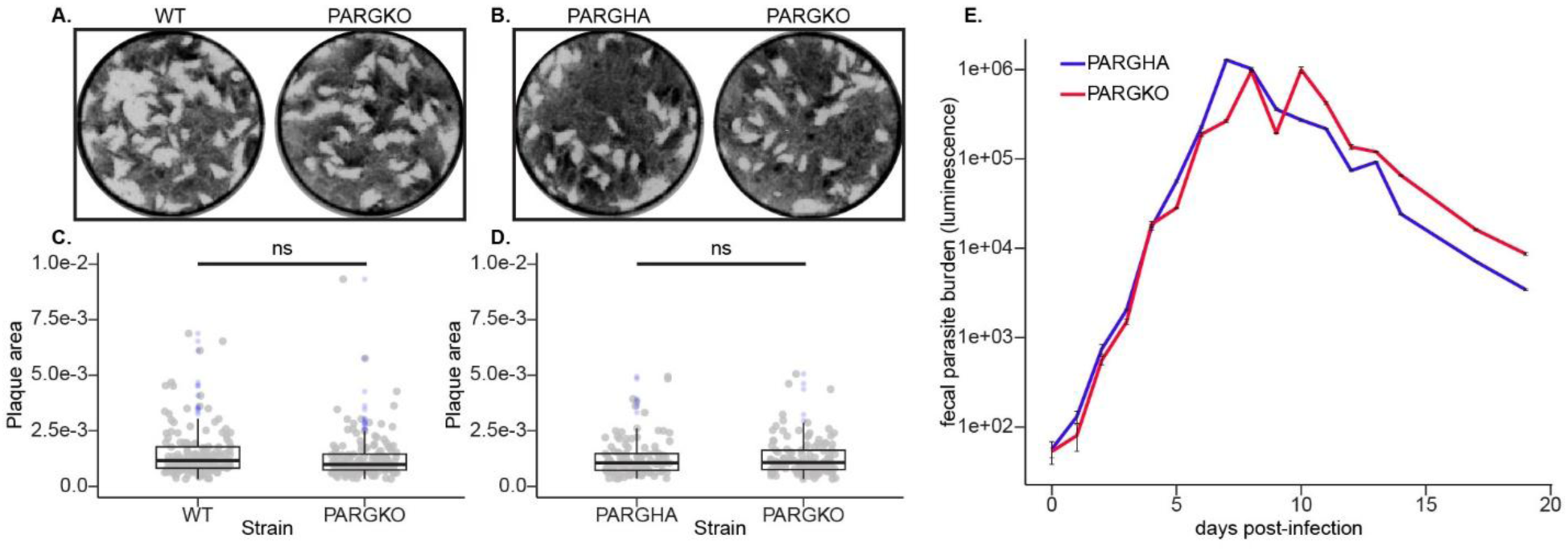
Dense granule PARG-domain containing proteins are not required for parasite fitness. **A-B.** HFF monolayers were infected with serial dilutions of the indicated *T. gondii*. Monolayers were fixed, stained with Giemsa, and imaged one week after infection. **C-D.** Plaque areas of three independent biological replicates per condition were segmented and their area measured. There was no significant difference in plaque area when comparing WT/TgPARG2-HA and TgPARG2-KO *T. gondii* strains. **E.** *Ifnγ^-/-^* mice were infected with passage-matched PARG-HA and PARG-KO *C. parvum* oocysts. Fecal luminescence was measured to assess parasite shedding over time. No difference in parasite shedding was observed in PARG-KO compared to PARG-HA *C. parvum.* Representative fecal luminescence curve from 3 independent experiments.

**Figure S4.**
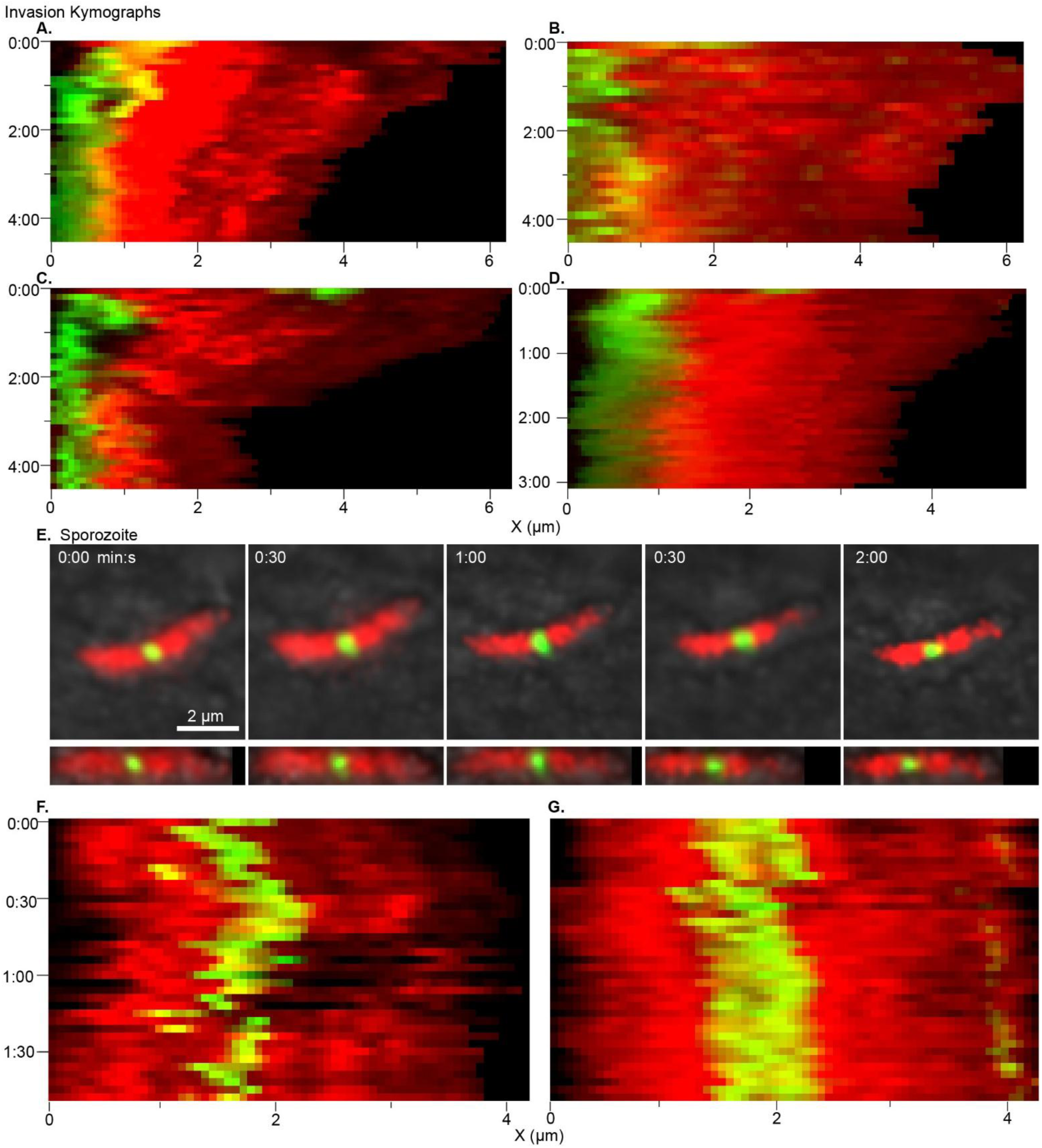
Additional time-lapse microscopy of PARG-mNeon secretion in invading and non-invading sporozoites. **A-D.** Additional kymographs showing the projected fluorescence intensity across the length of the parasite (X-axis) over time (Y-axis) from time-lapse microscopy of invading parasites showing changes in the distribution of PARG-mneon during invasion. The point of initial contact of the sporozoite with the HCT-8 cell is shown as time 0. **E.** Stills from a non-invading sporozoite imaged for 2 minutes. **F-G**. Kymographs showing the projected fluorescence intensity across the length of the parasite for two examples of non-invading sporozoites.

**Figure S5.**
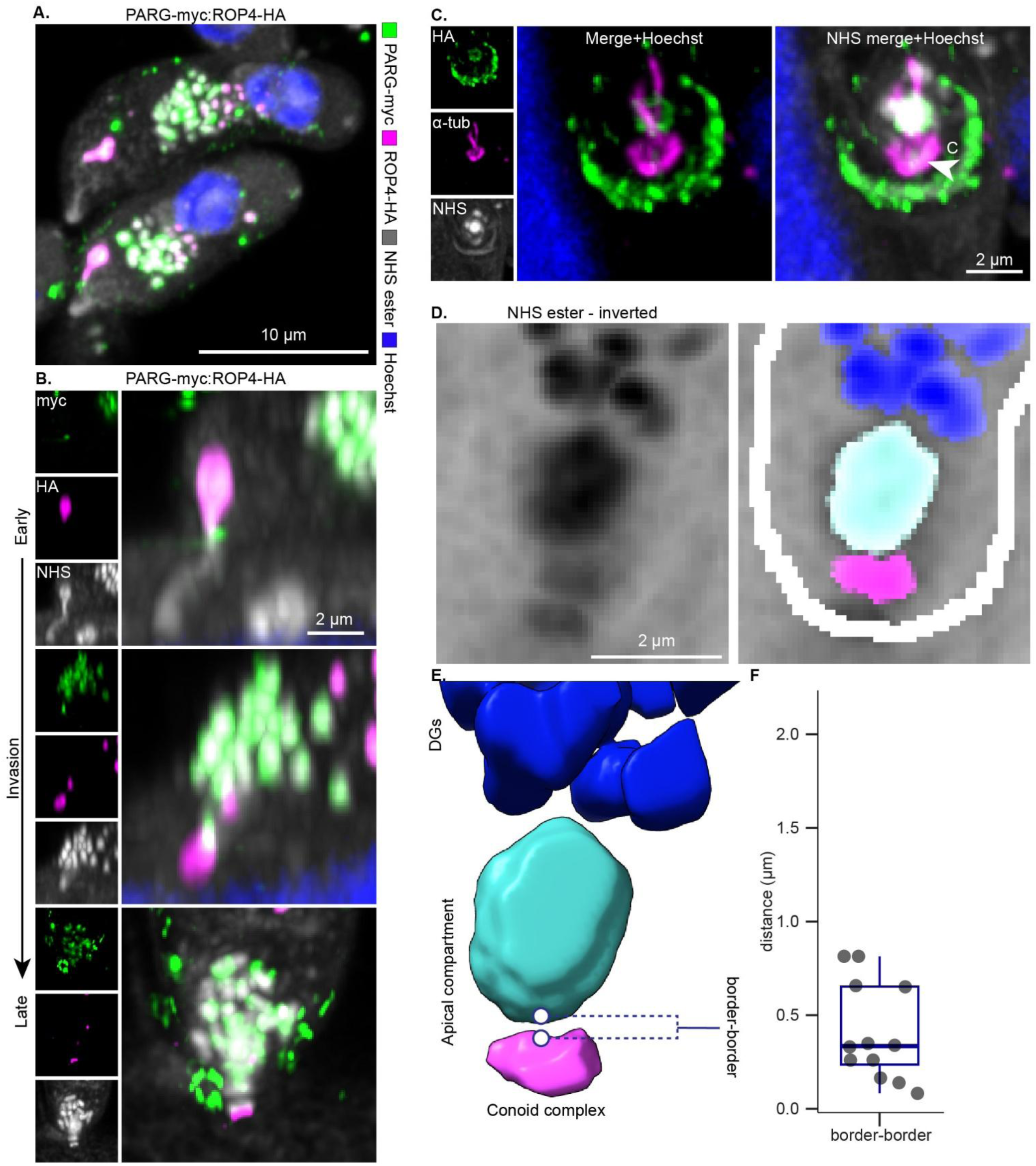
Rhoptry secretion precedes dense granule secretion, and the apical compartment is at a fixed position relative to the conoid complex. **A-B.** To simultaneously label the rhoptry and dense granule contents, PARG was myc-tagged at its endogenous location and Rop4-HA was knocked-in downstream with 500bp upstream of the start codon (PARG-myc:ROP4-HA). Myc is shown in green, HA is in magenta, NHS ester is in gray and Hoechst is in blue. **A.** merged panel showing two sporozoites labelled with ROP4 in the rhoptry bulb and PARG in the dense granules. **B.** The apical end of sporozoites at different stages of invasion are shown. The first example shows the rhoptry tip inserted into the host cell prior to discharge. In the next two panels, the rhoptry is no longer visible and the ROP4 signal has moved beyond the apical compartment. PARG-positive dense granules are still within the parasite. In the middle panel, they are beginning to cluster around the discharged rhoptry. In the last panel, they have started to fuse to form the apical compartment. **C.** Expansion microscopy of invading sporozoites expressing PARG-HA (staining in green) and stained with NHS ester (gray) and an α-tubulin antibody (magenta). The conoid complex (C) apical to the sub-pellicular microtubules is labelled with a filled arrow. Individual channels are shown on the left and tubulin-PARG-HA merge without and with NHS-ester are enlarged shown on the right. **D-F.** The apical compartment and the conoid complex were segmented in 3D using protein density from NHS ester labelling. **D.** An example of an inverted NHS ester contrast and segmentation overlay of the apical end of a parasite with the apical compartment (cyan), conoid complex (magenta), granules (blue), and parasite outline (white) shown. **E.** Rendering of the segmentation map in 3D. The distance measurement reported in F is illustrated. F. The shortest distance between the conoid complex and apical compartment borders was quantified and plotted on the right for 12 parasites. The edge of the apical compartment is 0.4 ± 0.26 µm (90 ± 47 nm scaled) away from the edge of the conoid complex.

**Fig. S6.**
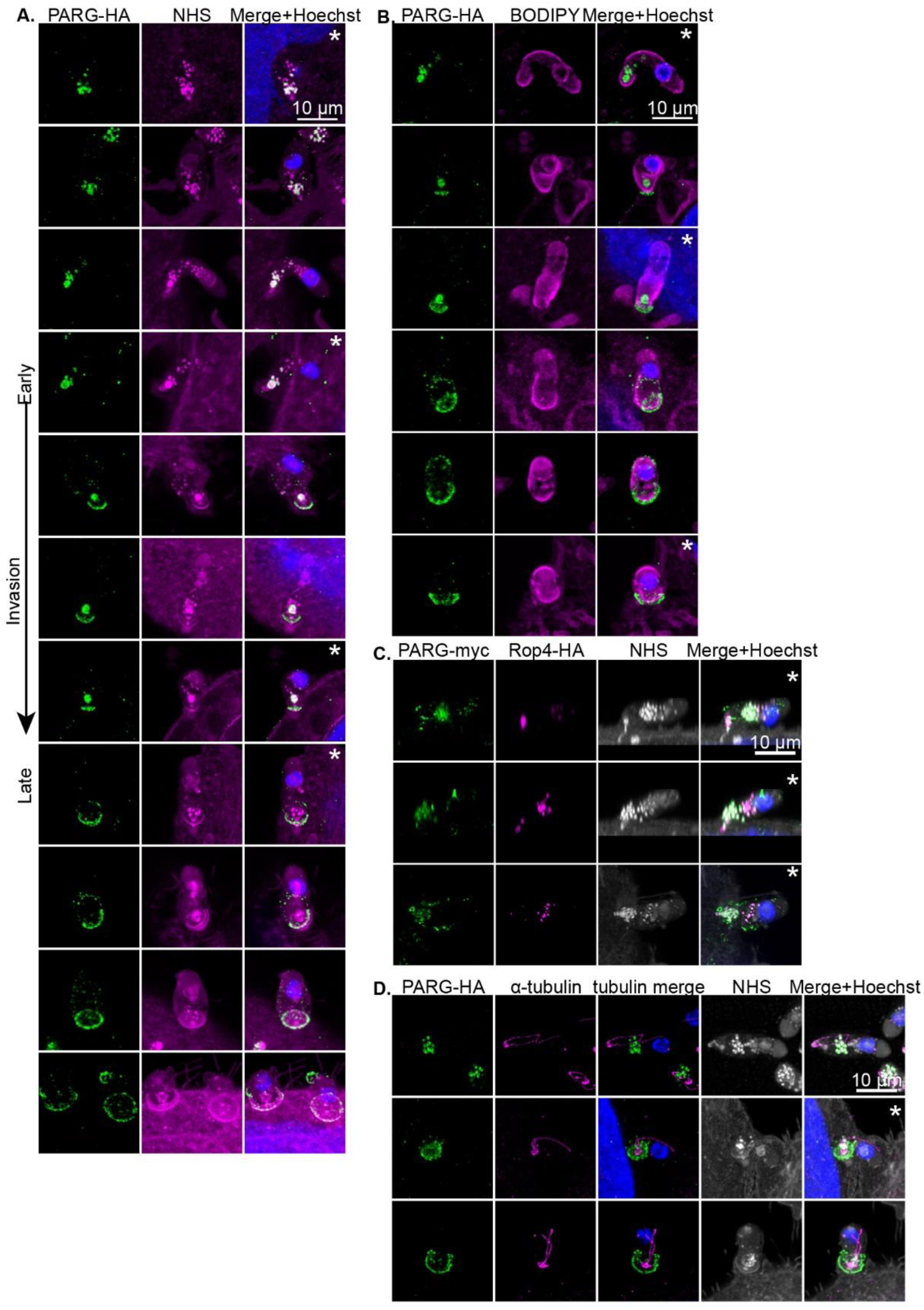
Expansion microscopy atlas of *C. parvum* invasion. **A-D.** Additional examples of invading parasites stained with the following markers: **A.** PARG-HA + NHS ester, **B.** PARG-HA + BODIPY, **C.** PARG-myc:Rop4-HA and **D.** NHS ester, PARG-HA and tubulin. Images marked with an asterisk are whole parasite views that are cropped in Figs 4 and S5.

**Fig. S7.**
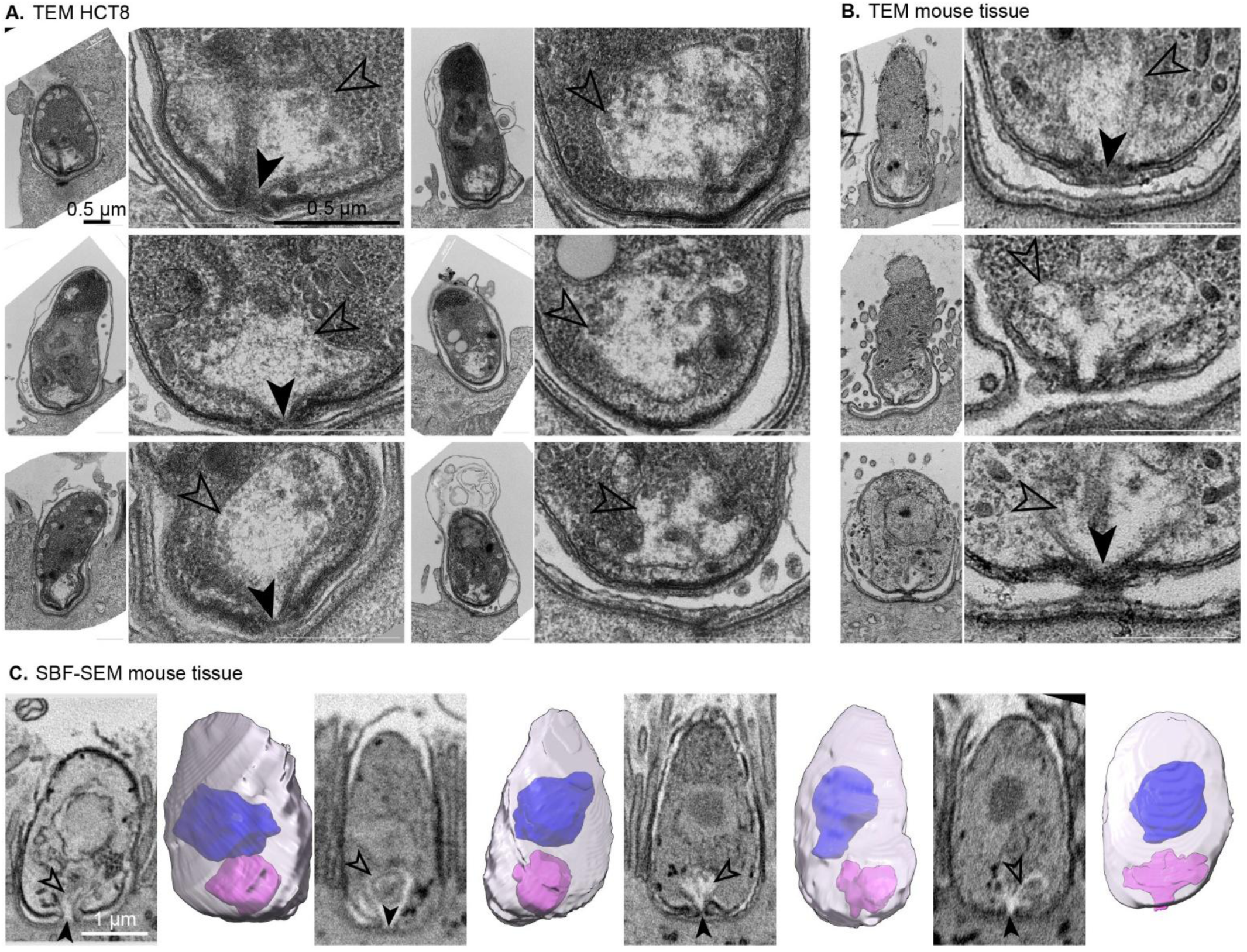
Electron microscopy atlas of *C. parvum* invasion. Additional thin section TEM images of invading sporozoites and immature intracellular trophozoites interacting with HCT-8 cells (**A.**) and from infected mouse intestines (**B)**. Details on the right-hand side show the apical end. An electron translucent, membrane-bound compartment can be seen in all examples. **C.** Selected central slices and segmentation of the parasite membrane (gray), nucleus (blue) and apical compartment compartment (magenta) from serial block-face SEM imaging of intestinal sections from infected mice. Where visible, the boundary/membrane of the apical compartment is marked with an open arrow and the conduit connecting the apical compartment to the host cytosol is labelled with a filled arrow.

**Figure S8.**
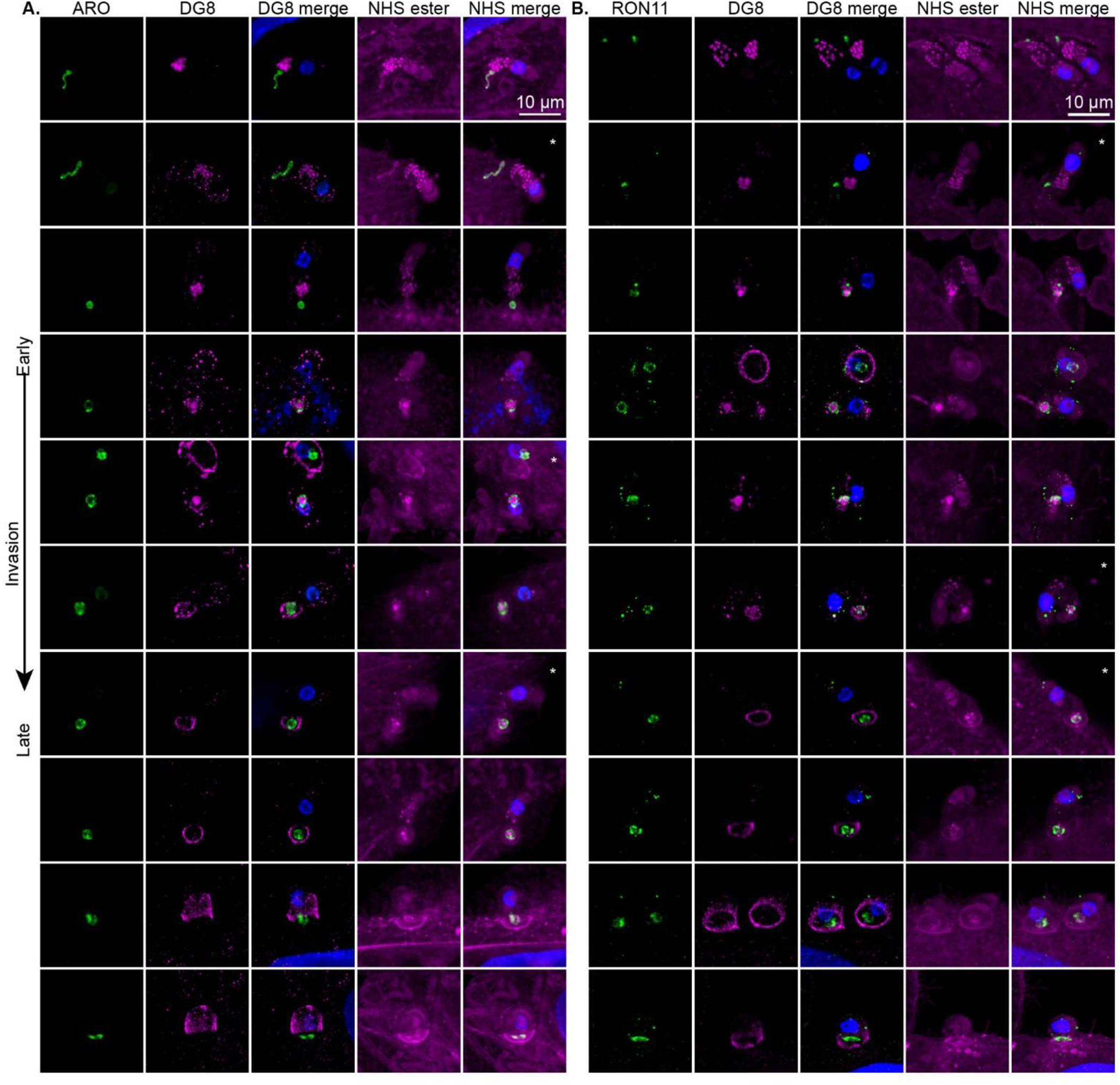
Expansion microscopy of rhoptry membrane markers during invasion. Additional examples of invading sporozoites and immature intracellular parasites visualized by expansion microscopy. Parasites expressed CpARO-HA (**A**) or CpRON11-HA (**B**). Immunofluorescence was used to detect HA epitopes (green) and DG8 (magenta). NHS ester was used to label protein density (Magenta) and Hoechst to label nuclei (blue). Images are arranged according to invasion stage from top to bottom. Images with an asterisk are full parasite views of cropped panels in Fig. 6.

**Figure S9.**
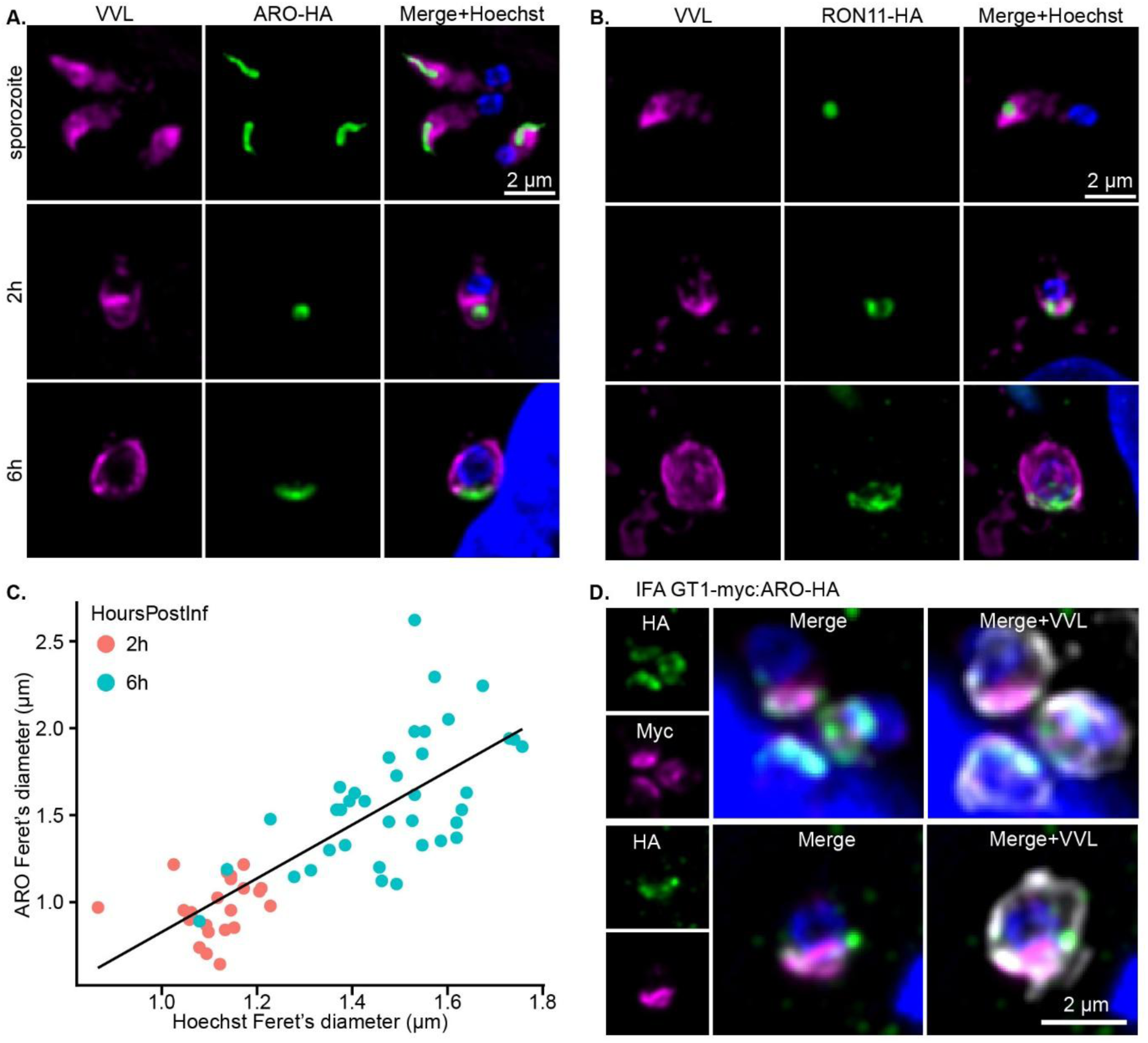
Immunofluorescence of CpARO and CpRON11 during trophozoite development. Representative examples of *C. parvum* parasites expressing **A.** CpARO-HA or **B.** CpRON11-HA at the sporozoite stage, the early trophozoite stage (2 hours post-infection) and the late trophozoite stage (6 hours post-infection. **C.** The size of the nucleus and ARO-HA-positive region were measured in maximum intensity projections. The maximum Feret’s diameter of the nucleus (Hoechst) and the ARO-HA region are plotted. Points represent a single parasite, colored according to time point at which they were fixed (red = 2 hours post-infection, teal = 6 hours post-infection). Trendline: Pearson’s Correlation Coefficient = 0.715, P-value = 1.38X10^-10^. Mean Feret’s Diameter of ARO at 2h = 0.96 ± 0.16 µm, at 6h = 1.59 ± 0.37 µm. **E.** Immunofluorecence of GT1-myc and CpARO-HA localization in mature 1N meronts. HA is green, Myc is in Magenta, VVL is in gray, and Hoechst is in blue.

**Figure S10.**
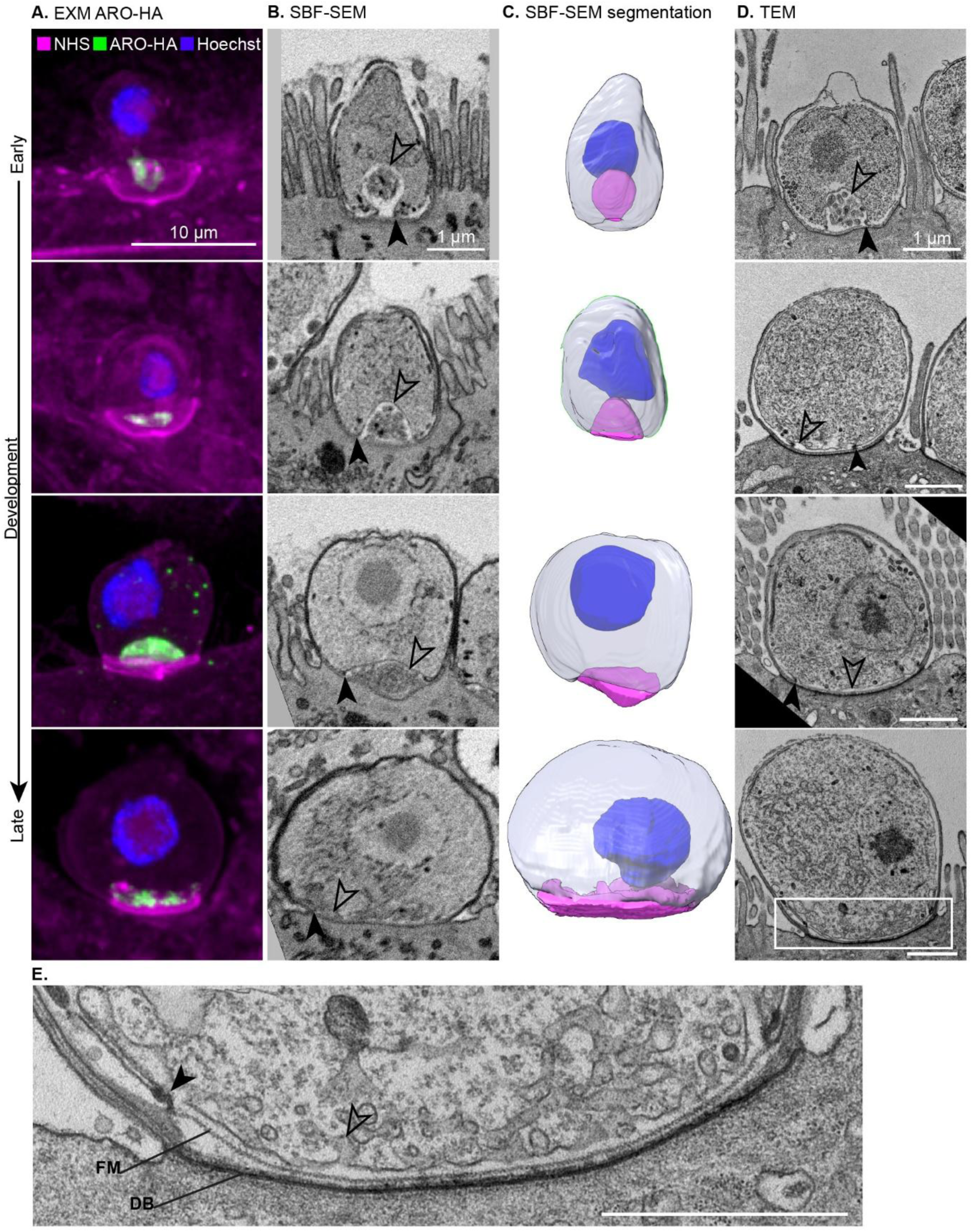
Comprehensive electron microscopy and expansion microscopy of feeder organelle development. **A.** Expansion microscopy and ARO-HA staining at progressive stages of parasite development. **B.** Central slices from selected SBF-SEM volumes from infected mouse intestines showing the apical compartment transitioning into the feeder organelle (top to bottom). The boundary of the apical compartment is shown with an open arrow and the annular ring with a filled arrow. **C.** Segmentation of the nucleus (blue), apical compartment/feeder organelle (magenta) and parasite boundary (transparent gray) and rendering in 3D. **D.** Thin section TEM from infected mouse intestines showing the apical compartment transitioning into the feeder organelle (top to bottom). The boundary of the apical compartment is shown with an open arrow and the annular ring with a filled arrow. **E.** Detail from thin section TEM (cropping area shown in the last image in panel **D)** showing the host-parasite interface of a mature intracellular parasite at high resolution. The dense band (DB), filamentous mesh (FM), feeder organelle membrane (open arrow) and annular ring junction (filled arrow) are labelled.

## Supplementary Tables

**Table S1.**
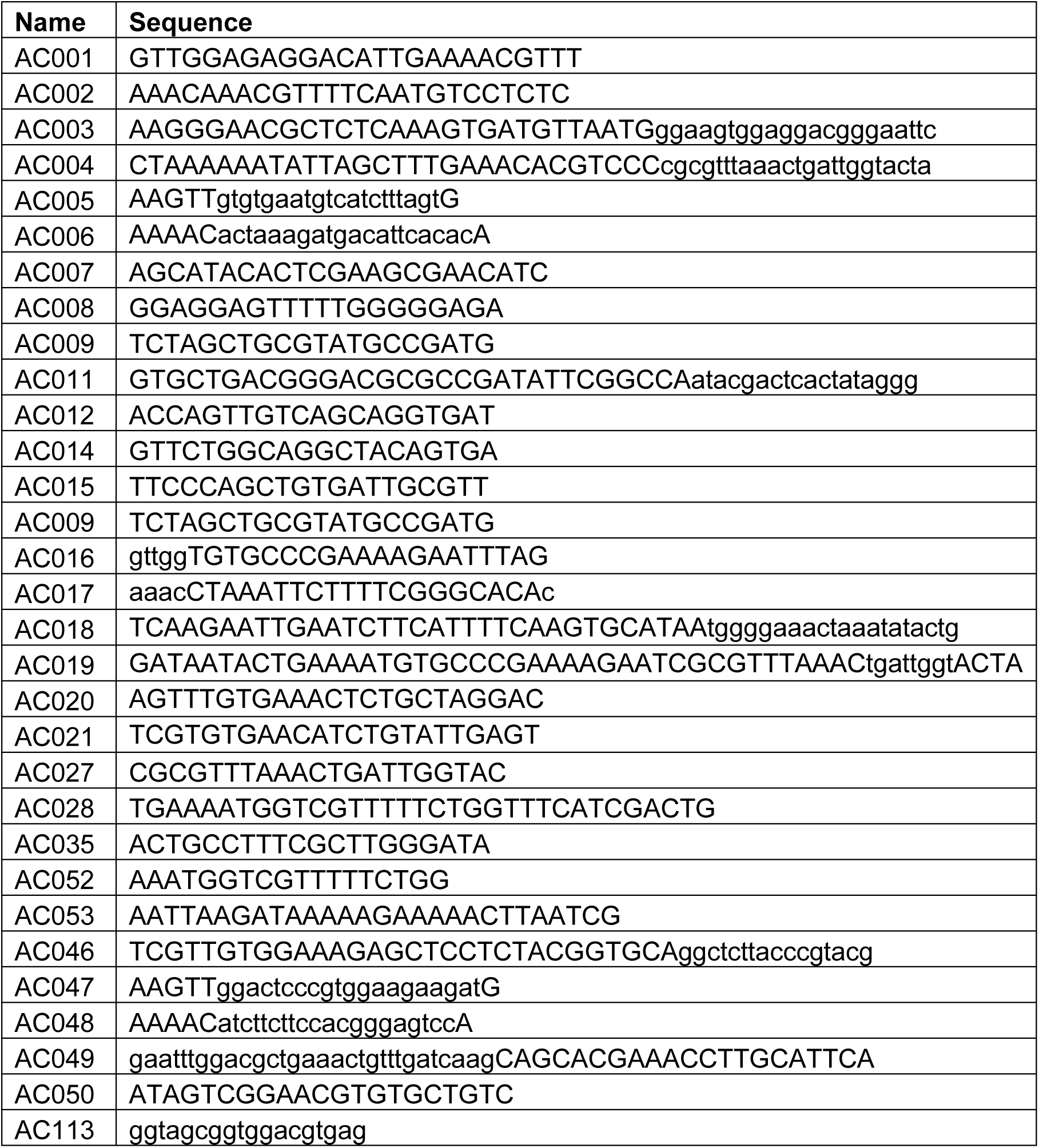
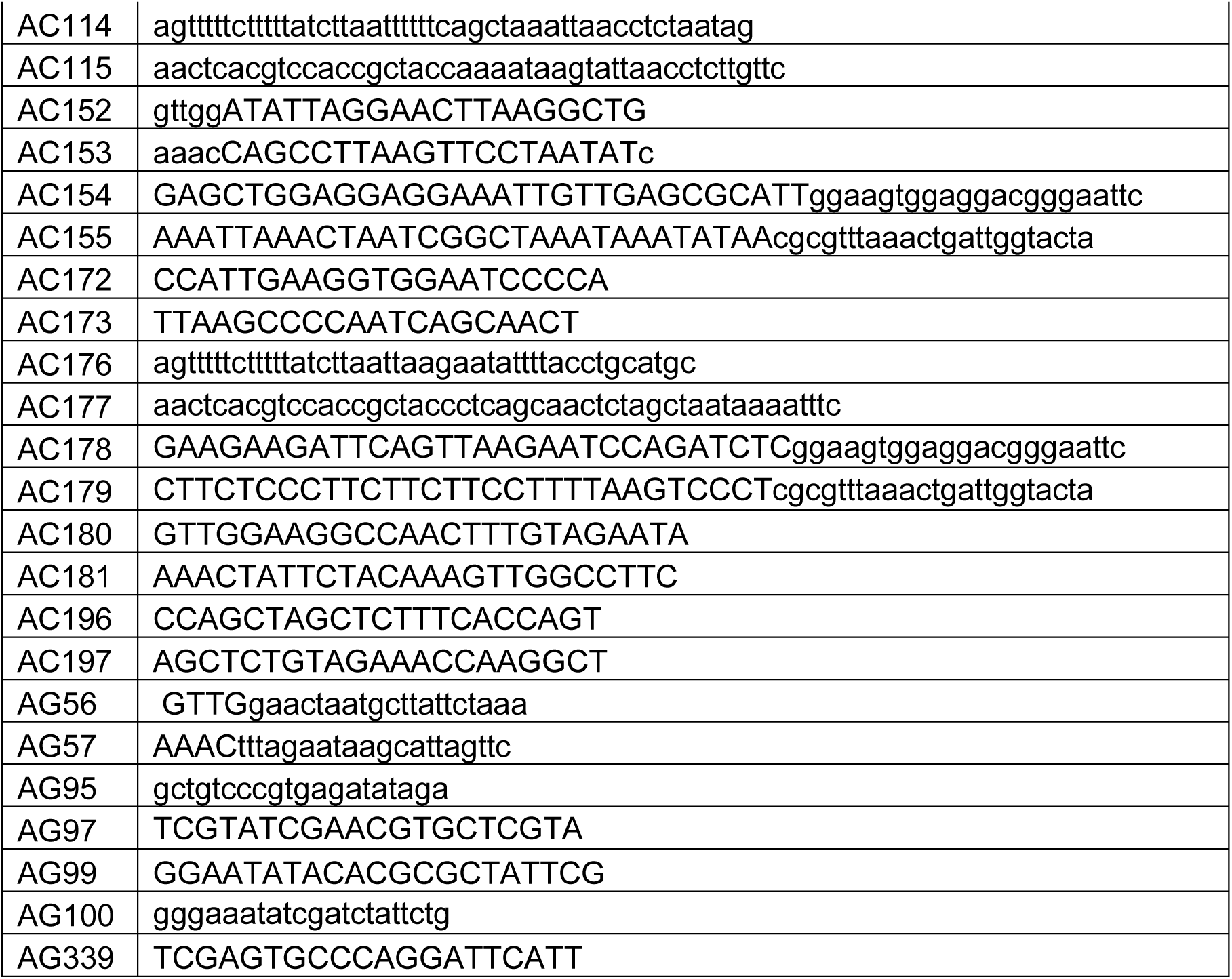
Oligonucleotides used in this study.

**Table S2.** Oligonucleotide pairs for guide RNA cloning and homology-directed repair template amplification.

| Transgenic | GeneID(s) | Guide oligos | repair oligos |
| --- | --- | --- | --- |
| PARG-HA | Cgd8_2160 | AC001/AC002 | AC003/AC004 |
| PARG-KO | Cgd8_2160 | AC016/AC017 | AC018/AC019 |
| PARG-mNeon:tdTomato | Cgd8_2160 | AC001/AC002 | AC003/AC004 |
| TgPARG2-HA | TgRH-280380 | AC005/AC006 | AC046/AC011 |
| TgPARG2-KO | TgRH-280380 | AC047/AC048 | AC049/AC011 |
| PARG-tdTomato | Cgd8_2160 | AC001/AC002 | AC003/AC004 |
| PARG-myc:Rop4-HA | Cgd8_2160,<br>Cgd3_1730 | AC001/AC002 | AC003/AC004 |
| CpARO-HA | Cgd2_370 | AG56/AG57 | AC142/AC143 |
| CpRON11-HA | Cgd3_2010 | AC152/AC153 | AC154/AC155 |
| CpGT1-myc:ARO-HA | Cgd3_4070,<br>Cgd2_370 | AC180/AC181 | AC178/AC179 |

**Table S3.**
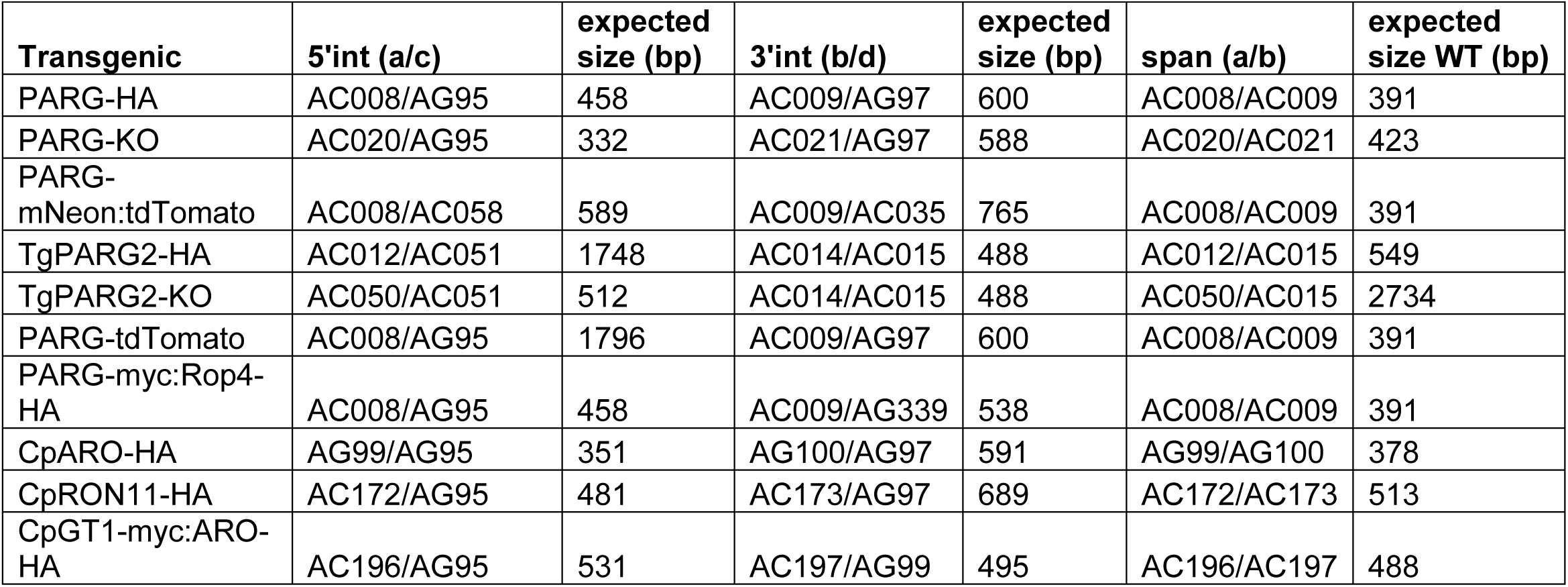
Oligonucleotide pairs for integration PCR and expected sizes.

| Transgenic | 5'int (a/c) | expected size (bp) | 3'int (b/d) | expected size (bp) | span (a/b) | expected size WT (bp) |
| --- | --- | --- | --- | --- | --- | --- |
| PARG-HA | AC008/AG95 | 458 | AC009/AG97 | 600 | AC008/AC009 | 391 |
| PARG-KO | AC020/AG95 | 332 | AC021/AG97 | 588 | AC020/AC021 | 423 |
| PARG-mNeon:tdTomato | AC008/AC058 | 589 | AC009/AC035 | 765 | AC008/AC009 | 391 |
| TgPARG2-HA | AC012/AC051 | 1748 | AC014/AC015 | 488 | AC012/AC015 | 549 |
| TgPARG2-KO | AC050/AC051 | 512 | AC014/AC015 | 488 | AC050/AC015 | 2734 |
| PARG-tdTomato | AC008/AG95 | 1796 | AC009/AG97 | 600 | AC008/AC009 | 391 |
| PARG-myc:Rop4-HA | AC008/AG95 | 458 | AC009/AG339 | 538 | AC008/AC009 | 391 |
| CpARO-HA | AG99/AG95 | 351 | AG100/AG97 | 591 | AG99/AG100 | 378 |
| CpRON11-HA | AC172/AG95 | 481 | AC173/AG97 | 689 | AC172/AC173 | 513 |
| CpGT1-myc:ARO-HA | AC196/AG95 | 531 | AC197/AG99 | 495 | AC196/AC197 | 488 |

## Supplementary Movie Legends

**Movie S1. PARG-mNeon:tdTomato invasion (related to Fig. 2, S4).** Time lapse microscopy of 5 invasion events. Sporozoites express PARG-mNeon (green) to label the contents of dense granules and cytosolic tdTomato (red). Bright field/DIC is shown in gray. Invasion movies 1-4 were acquired using an OMX SR Delta Vision microscope. Invasion movie 5 was recorded using a CrestOptics X-light V3 Spinning Disk Confocal. Additional details are available in the Method Details section. Movies are cropped to show frames prior to significant photobleaching. Scale bar = 5 µm.

**Movie S2. PARG-mNeon:tdTomato non-invading (related to Fig. 2, S4).** Time lapse microscopy of 3 attached sporozoites that do not invade. Sporozoites express PARG-mNeon (green) and cytosolic tdTomato (red). Brightfield is in gray. Images were acquired using an OMX SR Delta Vision microscope. Scale bar = 5 µm.

**Movie S3. PARG-tdTomato on Lifeact-GFP HCT-8 cells invasion (related to Fig. 3A-B).** Time lapse microscopy of a sporozoite expressing PARG-tdT (green) invading an HCT-8 cell expressing GFP-lifeact (red). Brightfield is in gray. Images were acquired using an OMX SR Delta Vision microscope. Scale bar = 2 µm.

**Movie S4. PARG-tdTomato on PLC-δ-GFP HCT-8 cells invasion (related to Fig. 3C-D)** Time lapse microscopy of a 2 invasion events. Sporozoites express PARG-tdT (green), and HCT-8 cell expressing GFP-PLC-δ (red). Brightfield is in gray. Image 1 was acquired using an OMX SR Delta Vision microscope. Image 2 was acquired using a CrestOptics X-light V3 Spinning Disk Confocal. The bright-field channel is not shown in Image 2 due to focus issues. Scale bar = 2 µm.

**Movie S5. Segmentation and rendering of expansion microscopy volume (related to Fig. 4, S5).** Animation through a Z stack of an invading parasite expressing PARG-HA imaged by expansion microscopy. PARG-HA is in green, NHS ester is in magenta and Hoechst is in blue. Amination through the Z stack showing just NHS ester displayed with inverted contrast (black on white). Segmentation of key features (parasite = gray, conoid complex = magenta, granules = blue, apical compartment = cyan) is overlayed in the next stack, and the 3D rendering is shown last. Scale bar = 10 µm.

**Movie S6. SBF-SEM volume of invading parasites, segmentation, and rendering (related to fig. 5C).** Amination through 5 SBF-SEM Z-stacks of invading (Volume 1) and early-stage intracellular (Volume 2-5) *C. parvum* parasites in mouse intestinal tissue. Segmentation and rendering of the apical compartment (magenta), parasite (gray) and nucleus (blue) are shown. Scale bar = 0.5 µm.

**Movie S7. TEM tomography of invading parasites (related to fig. 5E).** Amination through a tomogram in Z reconstructed from a tilt-series acquired from a *C. parvum* sporozoite in the process of invading an HCT-8 cell *in vitro*. Scale bar = 0.5 µm.

**Movie S8. SBF-SEM volume and segmentation of immature, maturing, and immature intracellular parasites (related to Fig. S10).** Animation through 4 SBF-SEM Z stacks of parasite in mouse intestinal tissue with rendering of the apical compartment/feeder (magenta), parasite (gray), and nucleus (blue) segmentations. Volumes illustrate progressive development of the apical compartment into the feeder organelle as intracellular parasites mature. Scale bar = 1 µm.

## Materials and Methods

### Parasite strains

*Cryptosporidium parvum* Iowa II oocysts were purchased from Bunch Grass farms. Transgenic parasites were generated and propagated using *ifnγ^-/-^*mice bred in house as described above. Oocysts were purified from feces as previously described using sucrose flotation followed by a cesium chloride gradient[77]. Oocysts were stored at 4°C.

*T. gondii RH ΔKu80* was propagated in human foreskin fibroblasts (HFF-1, SCRC-1041).

### Host cell culture

HCT-8 cells were obtained from ATCC and cultured in DMEM supplemented with 10% cosmic calf serum (CCS) and 20 mM L-Glutamine.

HCT-8 cells stably expressing GFP-lifeact and GFP-PLC-δ were generated previously by lentiviral transduction and cloned by limiting dilution[11].

Human foreskin fibroblasts were obtained from ATCC and cultured in DMEM supplemented with 10% cosmic calf serum (CCS), 20mM L-glutamine, and 100 µg/mL penicillin/streptomycin.

### Plasmid generation

To generate a plasmid allowing for cytosolic tdTomato labelling downstream of mneon tagging, the pLic mneon plasmid was digested with ClaI and PshAI, and T2A tdTomato was amplified from the previously generated[18] pLic nLuc-Neo-T2A-tdTomato-H2B-mNeon plasmid using primers AC052/AC053. The two fragments were assembled using Gibson Assembly (NEB), and correct assembly was verified using PCR with primers spanning the fragments (AC027/AC028, AC027/AC035) and sequencing. To generate a plasmid to knock-in CpRop4-HA (Cgd3_1730) downstream of an endogenous myc epitope tag, the CpLic-myc-atub-DD-MybHA-OE vector generated previously[78] was amplified using primers AC113/AC053. Genomic DNA isolated from bunch grass parasites was used as a template to amplify the CpRop4 ORF and 450bp of upstream sequence using primers AC114/AC115. These fragments were used in Gibson assembly to generate the plasmid that was verified using PCR (AG339 and AC027) and sequencing. Similarly, to express ARO-HA downstream of an endogenous Myc tag, the CpLic-myc-atub-DD-MybHA-OE vector was amplified using primers AC113/AC053. The CpARO ORF with 400bp of upstream sequence was amplified using AC176/AC177. The plasmid was generated from these fragments using Gibson assembly, and validated using sequencing and PCR (AG99, AC055). Guide RNA plasmids for *C. parvum* were generated as previously described[77]. Briefly, annealed oligonucleotides designed using the CRISPR guide RNA design tool for eukaryotic pathogens[79] were ligated into the *C. parvum* Cas9/U6 plasmid. Guide RNA plasmids for *T. gondii* were generated as described previously[80]. Guide RNA oligos were annealed and ligated into the *T. gondii* Cas9 plasmid (kind gift of Sebastian Lourido) linearized with BsaI. Oligo pairs used for guide RNA cloning are listed in table S2. Oligo sequences are in table S1.

### Generation of transgenic *C. parvum*

The method used for generation of transgenic *C. parvum* parasites is described in detail elsewhere[18, 77, 81]. Briefly, parasites were excysted with a 45 min incubation in 0.01 M HCl at 37°C and 5% CO_2_ followed by an incubation in a mixture of 0.2 mM bile salts (sodium taurodeoxycholate) and 20 mM sodium bicarbonate in PBS for 1 hour at 37°C and 5% CO_2_. Excysted sporozoites were mixed with DNA and electroporated with an Amaxa 4D device and used to infect *ifnγ^-/-^* mice. Mice were pre-gavaged with 8% sodium bicarbonate to neutralize stomach acid and infected with parasites by oral gavage. One day after transfection, transgenics were selected with paromomycin (16 g/L) provided in the drinking water. Feces were collected during parasite shedding, and parasites were purified using sucrose floatation and cesium chloride gradient centrifugation as previously described[77]. DNA for transfection was prepared as previously described and included 50 µg of Cas9/U6 plasmid with a guide RNA targeting a gene of interest, and a repair fragment generated by PCR amplification of the indicated CpLic tagging plasmid harboring the epitope or fluorescence tag and the nluc-neo casette with 20-30bp homology arms from the gene of interest. Guide and repair template were mixed and concentrated by ethanol precipitation. Primer sequences are listed in table S1. Primer pairs used to clone guide RNA targeting sequences into the Cas9/U6 plasmid, to amplify the CpLic plasmid to generate the repair template, and to check integration are listed in table S2.

### Generation of transgenic *T. gondii*

HA epitope tagging and knockout by gene replacement were performed as previously described[80]. Briefly, guide RNA oligos (TgPARG2-HA tagging: AC005/AC006, TgPARG2KO: AC047/AC048, Tables S1) were annealed and ligated into the toxoplasma Cas9 universal plasmid (gift from Sebastian Lourido) that was digested with BsaI. Templates for homology-directed repair containing a DHFR-cassette were amplified from the AID-HA DHFR plasmid (gift from Dominique Soldati) using the following primers with 30bp overhangs: HA tagging - AC046/AC011, KO – AC049/AC011. 40 µg Cas9/Guide RNA plasmid and PCR product were mixed and concentrated by ethanol precipitation. Freshly lysed *T. gondii* was electroporated using an ECM 630 Square Wave Electroporator. *T. gondii* was grown in 3 µM pyrimethamine to select for DHFR expression and cloned by limiting dilution.

### *T. gondii* plaque assays

*T. gondii* plaque assays were performed as previously described[82]. Briefly, freshly lysed *T. gondii* tachyzoites were used to infect HFF monolayers grown in 24-well plates and allowed to grow for 7 days. HFFs were then fixed with 4% PFA and stained with Giemsa stain and imaged using a BioRad GelDocXR+ with a white tray. Plaque size was quantified using basic thresholding and object analysis from the FIJI distribution of imageJ (v 1.54p)[83].

### Mouse infections

Animal protocols used were approved by the Institutional Animal Care and Use Committee of the University of Pennsylvania (protocol # 806292). *Ifnγ^-/-^* (stock no:002287) mice were purchased from Jackson Laboratory and bred in house in a mouse barrier facility. 5 to 8-week-old mice were used. All mice were sex and age matched within individual experiments. Both male and female mice were used, and no differences were observed. To passage transgenic parasites, *ifnγ^-/-^* mice were infected with 1X10^4^ oocysts in PBS by oral gavage. Paromomycin was included in the drinking water for the first three passages. To compare the relative fitness of PARG-HA and PARG-KO *C. parvum* parasites, age and sex-matched mice were infected with 1X10^4^ oocysts/mouse. Three biological replicates were performed. Fecal parasite burden was measured by nanoluciferase assay every other day as previously described[81].

### Immunofluorescence microscopy

HCT-8 or HFF cells were seeded onto 12 mm circumference #1.5 coverslips in 24-well plates. Oocyst excystation was triggered with a 5 min incubation in 25% sodium hypochlorite (bleach) and a 10 min incubation in 0.8% sodium taurodeoxycholate (bile salts) in PBS. Triggered oocysts were added to HCT-8 monolayers in DMEM + 1% CCS, pen/strep, and L-glutamine, and incubated at 37°C with 5% CO_2_ for the indicated amount of time. For *T. gondii*, HFF coverslips were infected with freshly lysed tachyzoites and incubated for 24h. Immunofluorescence was performed as previously described[18]. Coverslips were fixed with 4% paraformaldehyde (PFA) for 15 min at room temperature. To detect intracellular protein, coverslips were treated with 0.5% Triton X100 for 15 minutes at room temperature. Samples were blocked in 4% BSA (in PBS) for 1 hour and stained with primary antibodies in 4% BSA overnight at 4°C, washed 3 times with PBS, and incubated with secondary antibodies for 1 h at room temperature. Finally, Hoechst staining was performed for 5 minutes followed by 3 PBS washes, mounting on slides using Vectashield, and sealing with nail polish. For VVL protection assays, the permeabilization step was omitted and fluorescently labelled parasites interacting with HCT-8 cells were blocked with 4% BSA for 30 min, stained with VVL-biotin, washed 3 times in PBS, and stained with A647-conjugated streptavidin followed by hoechst staining, washing, and mounting.

For *in vivo* immunofluorescence, infected mice were euthanized and their small intestines were harvested and ‘swiss-rolled’ as previously described[84, 85]. Intestines were fixed overnight in 10% formalin at 4°C and incubated overnight in 30% sucrose prior to mounting in OCT. Cryo-sectioning was performed by the Penn Vet Comparative Pathology core facility (RRID: SCR 0022438). Immunofluorescence was performed as previously described[11, 21, 84, 85].

Samples were imaged using microscopes housed in the Penn Vet imaging core (RRID: L SCR_022764) using a Widefield Leica DM600B with a 100X (1.4 NA) oil immersion objective and deconvolved using Huygens deconvolution software (Scientific Volume Imaging, Hilversum, Netherlands). Images were also acquired on a Leica Stellaris 8 FALCON with a 100X (1.4NA) oil immersion objective. LIGHTNING processing was performed in LAS X software (Leica Microsystems Inc., Deerfield, IL). Optical sectioning was performed with a step size of 0.4-0.6 µm, and unless indicated otherwise, images represent maximum intensity projections of multiple Z-slices. Images were projected, rotated, and cropped using FIJI. The QuickFigures FIJI plugin was used to prepare and arrange split and merged channel views[86].

Quantification of the size of the region positive for CpARO staining and the size of the nucleus on maximum intensity projections was performed using FIJI. Channels were split and the hoechst and CpARO-HA channel were subjected to basic thresholding and object analysis to measure the area and Feret’s diameter (maximum diameter) of the indicated objects. The Pearson’s correlation coefficient was calculated using R (V4.5.3, The R Project for Statistical Computing) in Rstudio (V2023.06.1, Posit Software, PBC).

### Time-lapse microscopy of invasion

HCT-8 cells were seeded in 8-well CellVis chambered cover glass wells or ibidi 18-well glass-bottom imaging µ-slide wells. For 8-well chambers, 0.5-1X10^6^ oocysts were triggered as described above and added to the well. For 18-well slides, 2X10^5^ oocysts were added. Slides were mounted on an inverted OMX SR Delta Vision microscope with a 60X oil immersion objective (1.42 NA) equipped with a stage-top incubator maintaining temperature at 37°C and CO_2_ at 5%. For one experiment in which PARG-tdTomato parasites were imaged on PLC-GFP HCT-8 cells, samples were observed using a CrestOptics X-light V3 Spinning Disk Confocal housed in the Cell & Developmental Biology (CBD) Microscopy Core (RRID SCR_022373) with a 100X/1.45 NA oil immersion objective enclosed by an Okolab incubator. To minimize photobleaching, Samples were observed sporadically by DIC imaging until sporozoites were visible in the field of view, typically 45 minutes-to-1 hour after triggering and beginning incubation at 37°C. Once excysted sporozoites were visible, multi-channel Z-stacks (8-12 µm, 0.4 µm step sizes) were recorded every 3-5 seconds for 10-15 minutes, when photobleaching became severe. OMX stacks were aligned and deconvolved using Cytiva SoftWORx. Cropping, maximum intensity projection calculation, bleach correction, and denoising were performed in FIJI. For kymograph analysis, a segmented line was drawn along the length of the parasite in FIJI. The “straighten” tool was used with a line width of 20 pixels. The slicing tool was used to reslice the image along the X/T axis, and the resulting image was subjected to maximum intensity projection along the Y axis. Movies were selected in which the parasite was oriented perpendicular to the imaging axis for most of the movie.

### Ultrastructure expansion microscopy

Ultrastructure expansion microscopy (U-ExM) was performed as described previously[21, 63, 87]. Oocysts (0.5-1X10^6^) were triggered to excyst and added to HCT-8 cells seeded onto 12 mm coverglass circles in 24-well plates. After 2 hours, 6 hours, or 24 hours, coverslips were fixed with 4% PFA for 15 minutes followed by 3 PBS washes. Coverslips were incubated overnight at 37 °C in 1.4% formaldehyde/2% acrylamide in PBS. Gelation was performed by inverting coverslips on a 35 µl droplet of monomer solution (18% sodium acrylate, 10% acrylamide, 0.1% N,N’-methylene bisacrylamide in PBS) to which 5% tetramethylethylendiamine and 5% ammonium persulfate were added. Samples were incubated on ice for 5 min followed by 1h incubation at 37°C in a moist chamber. Gels were denatured at 95°C for 90 minutes in denaturation buffer (200 mM SDS, 200 mM NaCl, 50mM Tris pH9) followed by 3 30 min incubations in dH_2_O to expand the gels. At this stage, the diameter of the gels was measured to estimate the expansion factor. Gels were then incubated twice for 15 min in PBS, blocked in 4% BSA, and stained overnight at room temperature in primary antibody prepared in 4% BSA. Gels were washed 3 times for 10 min with PBS+0.2% Tween-20 and incubated in secondary antibody mixture, including NHS ester and hoechst in PBS, for 2.5 h before washing again in PBS+0.2% Tween-20 and re-expanded in dH_2_O. BODIPY staining was performed on expanded gels in 0.2% propyl gallate overnight at room temperature. Gels were mounted on 35mm glas-bottom mattek dishes coated with poly-D-lysine prior to imaging on a Stellaris 8 FALCON confocal microscope using a 63X water immersion objective (1.2 NA).

Leica LAS X software was used for LIGHTNING processing. Maximum intensity projections, image rotation, and cropping were performed in FIJI. The QuickFigures plugin was used as described above to generate split and merged views. To analyze the dimensions and distribution of the apical hub and conoid in invading parasites, thresholding and watershed separation were performed in FIJI on Z stacks to generate binary images. The 3D ROI manager (v4.1.7)[88] was used to segment objects in 3D and label different objects. The “Measure 3D” tool was used to measure the maximum diameter of the apical hub. The “Distances” tool was used to calculate the closest distance between the border of the apical hub and the border of the conoid in 3D.

### Transmission electron microscopy and electron tomography

HCT-8 cells grown on coverslips and infected with *Cryptosporidium parvum* were sequentially fixed with 4% paraformaldehyde in 0.05 M HEPES-buffered Hanks’ Balanced Salt Solution (HEPES-HBSS) for 1 h at room temperature, followed by 2.5% glutaraldehyde in 0.05 M HEPES-HBSS for at least 24 h. All subsequent preparation steps were carried out at room temperature. Post-fixation and block staining were performed sequentially with 1% osmium tetroxide, 0.1% tannic acid, and 2% uranyl acetate. Samples were dehydrated through an ascending ethanol series, infiltrated with ascending epoxy resin/acetone mixtures, and finally embedded in epoxy resin.

Sections were cut and examined using a transmission electron microscope (Tecnai Spirit, Thermo Fisher Scientific) equipped with a LaB6 filament and operated at 120 kV. For conventional TEM, 70 nm ultrathin sections were imaged with a side-mounted CCD camera (Phurona, EMSIS, Germany). For electron tomography, 150 nm thick sections were prepared and tilt series were acquired using a bottom-mounted Eagle camera (Thermo Fisher Scientific) and the TIA software with the integrated tomography module (Tecnai Imaging and Analysis, Thermo Fisher Scientific). Tomograms were reconstructed using the IMOD software package version 4.7[89] following the previously described procedure[90]. The apical compartment was manually segmented and measured in thin-section TEM images using ROI drawing and analysis tools in FIJI.

### Serial blockface SEM

Intestinal tissue from mice infected with *Cryptosporidium parvum* mutants was sequentially fixed with 4% paraformaldehyde in 0.05 M HEPES-HBSS for 1 h at room temperature, followed by 2.5% glutaraldehyde in 0.05 M HEPES-HBSS for at least 24 h. Subsequent preparation was followed as previously described[40] Briefly, an enhanced contrasting protocol including ferrocyanide-reduced osmium tetroxide post-fixation, thiocarbohydrazide-osmium liganding (ROTO), and a bloc staining with uranyl acetate and lead aspartate was performed. Samples were dehydrated through an ascending ethanol series, infiltrated with ascending epoxy resin/acetone mixtures, and embedded in epoxy resin.

SBF-SEM imaging and sectioning were performed using a field-emission scanning electron microscope equipped with an integrated ultramicrotome (Teneo Volumescope, Thermo Fisher Scientific). The microscope was operated at 2 kV and 100 pA in low vacuum (0.4 mbar) and images were recorded with the Volumescope Directional Backscatter (VS-DBS) detector at a pixel size of 4 nm (frame size 10,240 × 10,240 pixels) a dwell time of 1 µs per pixel and a section thickness of 40 nm. Image stacks were processed (slice registration, AI-based denoising[91], anisotropic diffusion filtering, and median filtering) using either Fiji[83] or Dragonfly 3D World (v2025.1, Comet Technologies Canada Inc.). Manual segmentation was performed in Amira 3D (v2024.2, Thermo Fisher Scientific) or Dragonfly 3D world (v2025.1, Comet Technologies Canada Inc.). Segmentation rendering was performed in UCSF Chimera X (V1.10)[92].

